# Novel mathematical morphology model identifies dorsal-ventral asymmetry of endothelial cell morphology in dorsal aorta of wild-type and Endoglin-deficient zebrafish embryos

**DOI:** 10.1101/2024.02.19.580931

**Authors:** Daniel Seeler, Nastasja Grdseloff, Claudia Jasmin Rödel, Charlotte Kloft, Salim Abdelilah-Seyfried, Wilhelm Huisinga

**Affiliations:** Institute of Mathematics, University of Potsdam, Germany; Institute of Biochemistry and Biology, University of Potsdam, Germany; PharMetrX Graduate Research Training Program: Pharmacometrics & Computational Disease Modelling, Berlin/Potsdam, Germany; Institute of Pharmacy, Freie Universität Berlin, Germany

**Author notes:** (https://www.pharmetrx.de/).

## Abstract

Endothelial cells, which line the lumen of blood vessels, locally sense and respond to blood flow. In response to altered blood flow dynamics during early embryonic development, these cells undergo shape changes that directly affect vessel geometry: In the dorsal aorta of zebrafish embryos, elongation of endothelial cells in the direction of flow between 48 and 72 hours post fertilization (hpf) reduces the vessel’s diameter. This remodeling process requires Endoglin; excessive endothelial cell growth in the protein’s absence results in vessel diameter increases. To understand how these changes in vessel geometry emerge from morphological changes of individual endothelial cells, we developed a novel mathematical model of the dorsal aorta’s apico-luminal surface that allows simultaneous quantification of vessel geometry and endothelial cell morphology. Based on fluorescently marked endothelial cell contours, we inferred cross-sections of the dorsal aorta that accounted for dorsal flattening of the vessel. By projection of endothelial cell contours onto the estimated cross-sections and subsequent triangulation, we finally reconstructed 3D surfaces of the individual cells. By simultaneously reconstructing vessel cross-sections and cell surfaces, we found that cell morphology varied between endothelial cells located in different sectors of the dorsal aorta in both wild-type and Endoglin-deficient zebrafish embryos: In wild-types, ventral endothelial cells were smaller and more elongated in flow direction than dorsal endothelial cells at both 48 hpf and 72 hpf. Although dorsal and ventral endothelial cells in Endoglin-deficient embryos had similar sizes at 48 hpf, dorsal endothelial cells were much larger at 72 hpf. In Endoglin-deficient embryos, elongation in flow direction increased between 48 hpf and 72 hpf in ventral endothelial cells but hardly changed in dorsal endothelial cells. Hereby, we provide evidence that dorsal endothelial cells contribute most to the disparate changes in dorsal aorta diameter in wild-type and Endoglin-deficient embryos between 48 hpf and 72 hpf.

**Author summary:** Endothelial cells, which form the innermost layer of each blood vessel, sense and respond to blood flow. During early embryonic development in zebrafish, endothelial cells of the dorsal aorta elongate in the direction of blood flow and hereby decrease the vessel’s diameter. To understand how these changes in vessel geometry emerge from morphological changes of individual endothelial cells, it is critical to precisely quantify both vessel geometry and cell morphology. To this end, we developed a 3D mathematical model of the dorsal aorta. Leveraging information from fluorescently marked endothelial cell contours allowed us to reconstruct the vessel’s surface. We applied this method to wild-type and mutant zebrafish embryos lacking functional Endoglin that is required for the physiological vessel diameter decrease. By quantifying vessel geometry and cell morphology in these embryos, we found that cell size and elongation in the direction of blood flow varied between endothelial cells located in different vessel sectors. Notably, we determined a subgroup of endothelial cells that contributed most to the vessel diameter increases in the absence of Endoglin. Future studies can investigate whether variability in endothelial cell behavior also contributes to the onset of human vascular malformations occurring due to a loss of Endoglin.

## Introduction

During embryonic development and adult life, the vasculature is continuously reshaped to maintain sufficient nutrient supply to tissues and to regulate systemic blood pressure. Endothelial cells (ECs), which line the lumen of blood vessels, can locally sense [1, 2] long-term changes of blood flow dynamics, occurring over a time span ranging between hours during embryogenesis [3] and weeks to months during adult life [4, 5]. As a consequence, vascular cells, i.e., ECs and mural cells, undergo behavioral changes that include alterations of cell shape, cellular rearrangements, migration, proliferation and cell death [6–8]. The combination of these cellular processes and structural alterations of the extracellular matrix results in the generation of new vessels via angiogenesis, the regression of weakly perfused vessels and changes in vessel luminal diameter; these coupled processes are called vascular remodeling [6–9]. It is crucial to improve our understanding of EC behavior during vascular remodeling as aberrations in sensing or responding to biochemical stimulation by blood flow on the level of ECs can cause diseases like atherosclerosis or a number of vascular anomalies [10–12].

Vascular remodeling can be precisely studied in zebrafish embryos, as their transparency and external development allows for non-invasive live-imaging of their vasculature over time. Additionally, many transgenic reporters are available that enable EC-specific fluorescent labeling of various cellular compartments [13]. The dorsal aorta (DA) is the major artery in the zebrafish trunk and undergoes drastic remodeling during embryonic development. Between 48 hours post fertilization (hpf) and 72 hpf, ECs in the DA elongate in the direction of blood flow [3]. As a consequence, the vessel length increases, but its diameter decreases. It was found that the TGF-beta co-receptor Endoglin is necessary for this remodeling process: Although ECs in Endoglin-deficient embryos also elongate in the direction of flow between 48 hpf and 72 hpf, excessive EC growth in the protein’s absence causes an increase in DA diameter instead [3].

To better understand the interplay of vessel geometry and cell morphology during vascular remodeling of the DA, it is critical to precisely quantify both DA geometry and EC surface morphology in the same embryo. While quantities like vessel diameter and EC perimeter can be directly measured in 3D microscopy images, mathematical models are required for the computation of other morphometric measures like the surface area of ECs, which surround the blood vessel. To learn how the disparate changes in DA diameter in wild-type and Endoglin-deficient zebrafish embryos emerge from EC shape changes, Sugden et al. computed 2D representations of ECs from their fluorescently labeled 3D cell contours [3]. For each EC *individually*, they first estimated an elliptic or hyperbolic cylinder from its contour. They then projected the EC contour onto the estimated cylinder and unrolled it. Hereby, the authors were able to quantify morphology of the unrolled ECs in 2D. This method, however, is not designed to compute an accurate description of vessel geometry: Cylinders estimated from different ECs whose surfaces contribute to the same vessel cross-section can differ substantially. Furthermore, by computation of a single cylinder per EC contour, variation in cross-sectional shape along the length of the cell is neglected.

To analyze the environment of ECs undergoing endothelial-to-hematopoietic transition in the DA, Lancino et al. computed a 2D representation of the DA from fluorescently labeled cell contours [14]. Along the length of a DA segment, they estimated circular cross-sections with variable centers but fixed radius that best fit the local fluorescent cell contour signal. To further account for variation in cross-sectional shape along the vessel axis, points on estimated circles were assigned the locally maximal fluorescence intensities in radial direction. The maximal fluorescence intensities were then unrolled onto a rectangle whose width was given by the DA’s length and its height by the cross-sectional circumference. While the unrolled map enabled the authors to visualize and quantify cell connectivity, it is not suited for precise analysis of either vessel geometry or cell surface morphology: By projecting fluorescence intensities onto a circle with fixed radius, variations in cross-sectional geometry in radial direction are dismissed. Furthermore, by changing the centers of the DA’s cross-sections along the vessel axis without incorporating these translations in the planar representation, cell contours become distorted. Quantification of morphology from complex cell contours can thus result in drastic deviations, e.g., in cell perimeter.

Finally, to study the spatial distribution of ECs within DA cross-sections, Campinho et al. reconstructed the 3D apico-luminal surface of the DA from fluorescent endothelial cytoskeleton markers [15, 16]. Along the DA’s length, they estimated elliptic cross-sections from local endothelial cytoskeletal signals. They then projected the centers of fluorescently labeled EC nuclei onto the reconstructed tube. While this method allowed the authors to quantify both the luminal area of the DA along its axis and the distribution of EC nuclei in different cross-sectional sectors of the DA, the authors did not leverage information from EC contours. Without this information, the boundaries of EC surfaces on the estimated vessel surface cannot be defined and thus, the individual cells’ surface morphology cannot be evaluated. While all the aforementioned models described DA cross-sections using circles or ellipses [3, 14, 15], we repeatedly observed in 3D microscopy images that dorsal segments of DA cross-sections can be flattened in comparison to their ventral segments.

To simultaneously and physiologically accurately describe cross-sectional geometry and EC surface morphology in the DA, we developed a novel mathematical morphology model of the apico-luminal surface of the vessel’s EC layer. By estimating both vessel cross-sections and EC surfaces solely from EC contours, our model guarantees consistency of vessel geometry and EC surface morphology. To account for the observed dorsal flattening of the DA, we developed a novel parametric cross-sectional shape model with inherent dorsal-ventral asymmetry. Following vessel surface reconstruction, cross-sectional geometry of the DA and EC surface morphology can be quantified within the same embryo. By quantification of EC surface morphology in 3D, we avoid cartographic distortions that would be caused by unrolling of the changing cross-sections along the vessel axis. Our preprocessing and surface reconstruction methods require little user intervention and thus enable statistical analyses of cross-sectional geometry and EC surface morphology over large numbers of embryos.

Using our mathematical model, we quantified DA geometry and EC surface morphology in wild-type and Endoglin-deficient zebrafish embryos at 48 hpf and 72 hpf and found good agreement with the results reported in [3]. Due to the precise reconstruction of cross-sectional geometry and EC surface morphology in the same blood vessels, our method further enabled us to identify a previously unrecognized dorsal-ventral asymmetry of EC surface morphology in the dorsal aorta: In wild-type embryos, ventral ECs were smaller and more elongated in the direction of blood flow than dorsal ECs at both 48 hpf and 72 hpf. Although dorsal and ventral ECs in Endoglin-deficient embryos had similar sizes at 48 hpf, dorsal ECs were much larger at 72 hpf. In Endoglin-deficient embryos, elongation in the direction of flow increased between 48 hpf and 72 hpf in ventral ECs but hardly changed in dorsal ECs. Hereby, we provide evidence that dorsal ECs contribute most to the disparate changes in dorsal aorta diameter in wild-type and Endoglin-deficient embryos between 48 hpf and 72 hpf.

## Materials and methods

### Data acquisition

We reconstructed DA geometry and EC surface morphology from manually annotated EC contours in 7 wild-type (wt) and 6 Endoglin-deficient (Eng-def) zebrafish embryos at 48 hpf and 72 hpf (see Table 1, analysis data). For model validation, we acquired independent data consisting of two wild-type embryos at 72 hpf where the vessel lumen was visualized by angiography (see Table 1, validation data). To allow quantification of annotation uncertainty, we annotated each EC contour twice in both of these embryos.

**Table 1.** Data used for the analysis of dorsal aorta geometry and endothelial cell morphology and for model validation.

| Data | Purpose | Phenotype | Time point | Embryo IDs | EC contours | Angiography |
| --- | --- | --- | --- | --- | --- | --- |
| analysis data | geometric and morphometric analysis | wt<sup>†</sup> | 48 hpf, 72 hpf | 1–7 | annotated once | no |
|  |  | Eng-def<sup>‡</sup> |  | 8–13 |  |  |
| validation data | model validation | wt | 72 hpf | 14–15 | annotated twice | yes |

<sup>†</sup>: Wild-type (wt) embryos carry maximally one *eng*<sup>*mu130*</sup> loss-of-function allele. <sup>‡</sup>: Endoglin-deficient (Eng-def) embryos carry two *eng*<sup>*mu130*</sup> loss-of-function alleles.

### Zebrafish genetics and maintenance

Handling of zebrafish was done in compliance with German and Brandenburg state law, carefully monitored by the local authority for animal protection (LAVG, Brandenburg, Germany) and in full accordance with the European Directive 2010/63/UE regarding the protection of animals used for scientific purposes. The following strains of zebrafish were maintained under standard conditions as previously described [17]: Zebrafish with the transgenic reporter *Tg(fli1:pecam1-EGFP)*^*ncv27*^ express the fusion protein Pecam1-EGFP specifically in ECs due to the *fli1* promoter; fusion of Pecam1 to EGFP fluorescently marks endothelial cell-cell junctions [18]. In zebrafish with the transgenic reporter *Tg(fli1:NLS-mCherry)*^*ubs10*^, nuclear localization of the fusion protein NLS-mCherry fluorescently marks EC nuclei [19]. Finally, *eng*^*mu130*^ zebrafish contain a frameshift mutation in the *eng* gene that results in a premature stop codon; nonsense-mediated decay of *eng*^*mu130*^ mRNA leads to a loss of function of the encoded Endoglin protein [3].

To allow identification of EC contours, all the zebrafish analyzed in this article expressed at least one allele of *Tg(fli1:pecam1-EGFP)*^*ncv27*^. Co-expression of one or two alleles of *Tg(fli1:NLS-mCherry)*^*ubs10*^ in a subset of the analyzed zebrafish further aided in the visual separation of neighboring ECs. Due to the recessive phenotype of the *eng*^*mu130*^ allele [3], we refer to embryos homozygous for the loss-of-function mutation *eng*^*mu130*^ as Endoglin-deficient embryos and to all other embryos as wild-types.

### Microangiography and live imaging

For live imaging experiments, embryos were collected, incubated at 28.5 ^°^C and treated at 24 hpf with 1-phenyl-2-thiourea (PTU) (Sigma Aldrich) to inhibit pigmentation. Embryos were manually dechorionated, anesthetized by incubation in egg water containing 0.003 % Tricaine and embedded laterally in 1 % low melting agarose (Lonza 50081) containing 0.16 mg*/*mL Tricaine in glass-bottom dishes (MatTek) and imaged. For microangiography, anesthetized embryos were injected with 5 μg*/*μL Dextran TexasRed (70 kDa, D1864, Invitrogen) into the common cardinal vein at 48 hpf and immediately processed for imaging. Subsequently, embryos were recovered, incubated at 28.5 ^°^C in normal egg water, remounted at 72 hpf and imaged again. Imaging was conducted with an LSM 780 confocal microscope (Zeiss) using a 20x-objective. After imaging, all embryos were genotyped by PCR.

### General notation

Throughout the article, we denote the ordered sequence of 3D points of endothelial cell contour *i* = 1, 2, …, *N* that is the result of computational step *𝓁* by EC_*𝓁,i*_. For example, EC_anno,*i*_ is the manually annotated contour of endothelial cell *i*. Each sequence EC_*𝓁,i*_ consists of *n*_*𝓁,i*_ points:

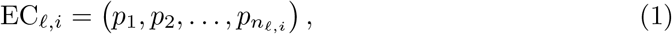

where we write the *x, y* and *z* components of *p* ∈ EC_*𝓁,i*_ as

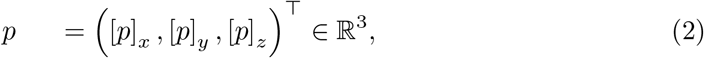

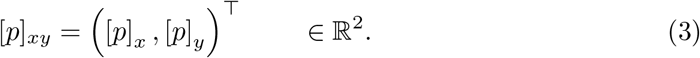

We denote the successor/predecessor of a point *p* ∈ EC_*𝓁,i*_ using *p*^+^ and *p*^−^, respectively:

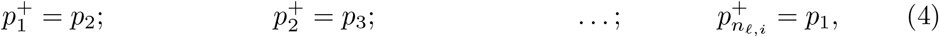

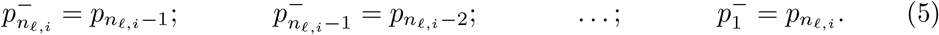

Concatenating the points of all cell contours (*i* = 1, 2, …, *N*), an ordered sequence

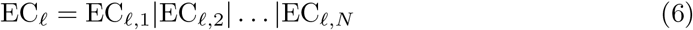

of length 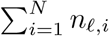 is obtained. Here, | denotes concatenation of tuples. Throughout the text, we use the multiplication sign both for the multiplication *α* · *v* of a scalar *α* ∈ ℝ and a vector *v* ∈ ℝ^3^ and for the inner product

*v*_1_ · *v*_2_ = [*v*_1_]_*x*_ · [*v*_2_]_*x*_ + [*v*_1_]_*y*_ · [*v*_2_]_*y*_ + [*v*_1_]_*z*_ · [*v*_2_]_*z*_ of two vectors *v*_1_, *v*_2_ ∈ ℝ^3^. The type of multiplication is clear from the dimensions of the factors.

We now explain the method for reconstruction and quantification of DA geometry and EC surface morphology from fluorescently marked ECs. An overview of the method is shown in Fig 1. To increase accessibility of the article, we provide a list of frequently used mathematical symbols in Table S1.

**Fig 1.**
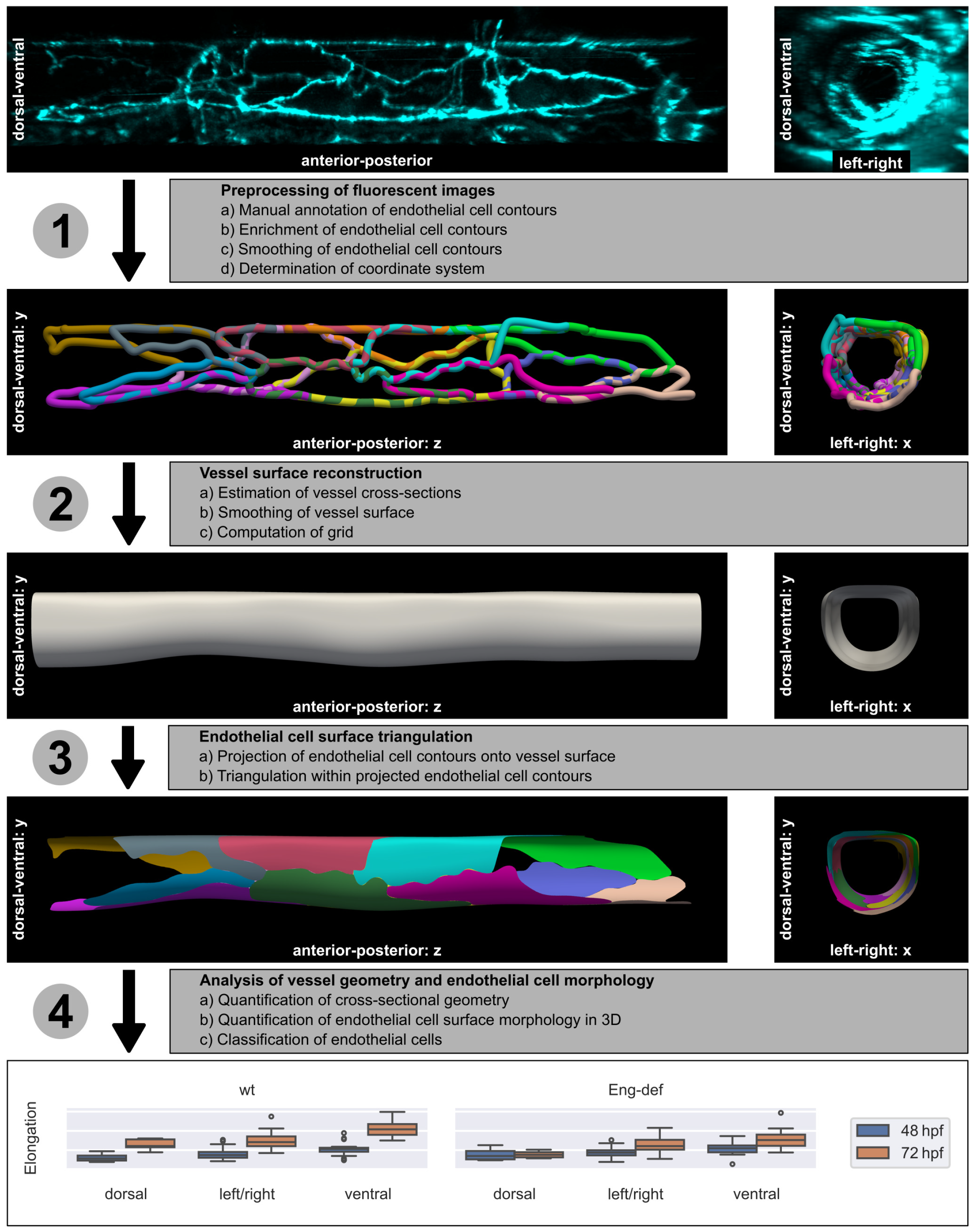
Method overview. The individual steps are explained in the text. Illustrations of EC contours and the DA’s surface are based on Endoglin-deficient embryo 13 at 72 hpf. Elongation of ECs was quantified using the analysis data (see Table 1).

### Step 1: Preprocessing of fluorescent images

#### a) Manual annotation of endothelial cell contours

To obtain EC contours within the DA, we employed Imaris Software V9.9.1 (Bitplane). Within a 3D reconstruction of the fluorescent signal from confocal z-stacks, we manually outlined Pecam1-EGFP-marked cell junctions over an area spanning 3–4 intersegmental vessels using the “Measurements” tool. During outlining, we excluded small ventrally located cells that were extruding from the vessel. We confirmed that points were placed in the center of the fluorescent signal by 3D image rotations. Each EC was annotated independently of its neighbors, i.e., joint cell-cell contacts were annotated twice. This increased the number of points available per cell-cell contact. We ensured that contour segments between consecutive points were well approximated by a straight line. This reduced the number of points needed to accurately describe EC contours. Additionally, we annotated vectors, defined as two ordered points, that provided approximate directions of anterior-posterior, dorsal-ventral and left-right axes. Data were exported as CSV files containing the points’ 3D coordinates. We denote the manually annotated contour of endothelial cell *i* as EC_anno,*𝓁*_.

For model validation against angiography, we additionally outlined cross-sections of the DA within 3D reconstructions of the fluorescent signal from confocal z-stacks of Dextran-perfused DAs. Using the “Ortho slicer” tool, we manually annotated the outlines of several cross-sections along the anterior-posterior axis of the DA with the “Measurements” tool. We confirmed that points of a cross-section lay approximately on a plane by 3D rotation. Again, we ensured that contour segments between consecutive points were well approximated by a straight line. In the remainder of the text, we refer to the outlined DA cross-sections as angiogram slices.

#### b) Enrichment of endothelial cell contours

As we annotated each EC independently of its neighbors, joint cell-cell contacts were annotated twice. To leverage this information in the inference process, we enriched each cell contour by points of its neighbors in an iterative process (see Fig 2A): We initialized the enriched cell contour EC_enri,*i*_ of cell *i* with its manually annotated cell contour EC_anno,*i*_. Then, we iterated over all points *q* ∈ EC_anno_ with *q* ∉ EC_anno,*i*_. Whenever a point *q* lay in at least one cylinder centered around an edge (*p, p*^+^) of the current version of the enriched contour EC_enri,*i*_, we inserted *q* into EC_enri,*i*_. To control which points were inserted, we defined two tuning parameters: the cylinder radius *r >* 0 and the length extension Δ*h >* 0 resulting in a cylinder length *h* = ||*p*^+^ − *p*||_2_ + 2Δ*h* for contour edge (*p, p*^+^).

**Fig 2.**
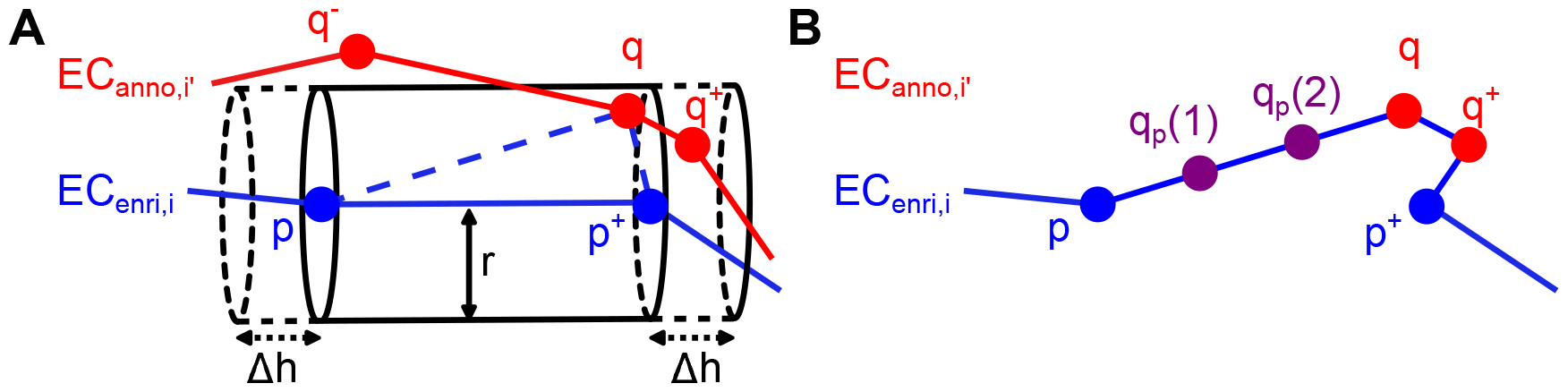
Cell contour enrichment. **(A)** Enrichment by neighboring cells. EC_enri,*i*_ is the current version of the enriched cell contour of cell *i*. EC_anno,*i*_^*′*^ is the manually annotated cell contour of cell *i*^′^. In the previous iteration, it was checked whether point *q*^−^ ∈ EC_anno,*i*_^*′*^ lies in any cylinder centered around an edge of EC_enri,*i*_. As *q*^−^ was too far away from these edges, it was not inserted into EC_enri,*i*_. In the current iteration, it is found that *q* ∈ EC_anno,*i*_^*′*^ lies in the cylinder centered around the edge (*p, p*^+^) of EC_enri,*i*_ with radius *r* and length *h* = ||*p*^+^− *p*|| _2_ + 2Δ*h*. Thus, *q* is inserted into EC_enri,*i*_. Following insertion of *q*, point *q*^+^ will be inserted into EC_enri,*i*_ in the subsequent iteration because it will be located within the cylinder centered around (*q, p*^+^). **(B)** Enrichment by interpolation. The enriched cell contour EC_enri,*i*_ resulting from adding both *q* and *q*^+^ is shown. Two points *q*_*p*_(1) and *q*_*p*_(2) obtained by linear interpolation are added on the edge (*p, q*).

Since we annotated fewer points in straight contour segments, the density of annotated points varied along contours. To homogenize this point density, we added *n*_interp_(*p*) additional points on each contour edge (*p, p*^+^) by linear interpolation (see Fig 2B). To ensure that the point density on all edges was comparable, we scaled the number *n*_interp_(*p*) of points interpolated on contour edge (*p, p*^+^) with the edge length:

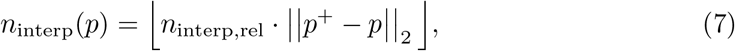

where *n*_interp,rel_ is a tuning parameter controlling the relative number of points interpolated per unit length (here: μm) on edges of any contour. We provide further details on the cell contour enrichment in Section 1 of S1 Appendix.

#### c) Smoothing of endothelial cell contours

To obtain a continuous representation of each cell contour, we employed cubic spline curves. Cubic splines are defined by piecewise polynomials of degree 3. These functions and their first and second derivatives are continuous and can thus accurately describe both concave and convex regions of cell contours. More specifically, we fitted periodic cubic smoothing splines [20]

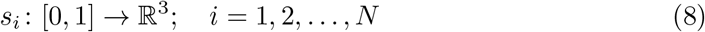

to the enriched cell contours. These curves minimize discontinuities in the third derivative while keeping the sum of squared distances between the points of the enriched contours and their projections onto the splines below or equal to a user-specified threshold *ϵ*_spline,*i*_ *>* 0. To ensure that splines fitted equally well to contours of small and large cells, we scaled the individual error thresholds *ϵ*_spline,*i*_ by the number *n*_enri,*i*_ of points on the respective enriched cell contours:

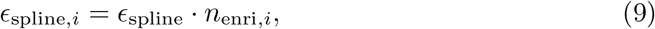

where *ϵ*_spline_ is a tuning parameter that serves as an upper bound on the mean squared distance for all splines. We parameterized each spline by the normalized arc length of the corresponding enriched cell contour.

Finally, we computed equidistant points on each endothelial cell contour spline. We denote the computed equidistant data points on the contour spline of cell *i* as

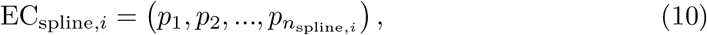

where

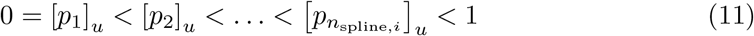

are the parameters of the points on the spline *s*_*i*_. To homogenize the density of points on EC contours, i.e., to take into account that large cells contain more information on the vessel surface than small cells, we scaled the number *n*_spline,*i*_ of points computed on each cell contour spline with the cell’s perimeter:

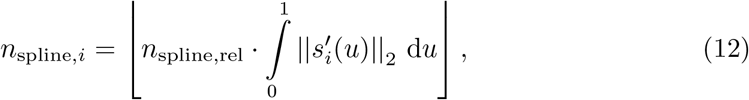

where *n*_spline,rel_ is a tuning parameter that controls the relative number of points computed per unit length (here: μm) for all splines. For more details on the computation of equidistant points on cell contour splines, see Section 5.1 of S1 Appendix.

#### d) Determination of coordinate system

The original coordinates of the cell contours depended on the position of the zebrafish embryo (and thus of the DA) within the mounting medium during imaging. As the DA is an approximately straight blood vessel without bifurcations, it is possible to define a straight line representing its anterior-posterior axis. Vessel cross-sections are then located on parallel planes perpendicular to this axis.

For each embryo at either time point, we defined the left-right (*x*), dorsal-ventral (*y*) and anterior-posterior (*z*) axes such that a single cross-sectional shape, termed mean shape, best fitted to all data points projected onto the *xy*-plane (see next section for the employed cross-sectional shape). The relationship between the coordinate system’s axes and the mean shape are illustrated in Fig S2. Our manually annotated vector for the dorsal-ventral axis was used to initialize the process of coordinate system determination. To ensure that the directionality of the anterior-posterior and left-right axis were not inverted, we compared the estimated axes with their respective manually annotated axis vectors. Below, the mean shape estimated together with the coordinate system is denoted 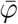 with parameters 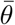. To differentiate the coordinates of contours in the original and the estimated coordinate system, we write *s*_transf,*i*_ for the spline of cell *i* within the new coordinate system and EC_transf,*i*_ for the equidistant points on the spline of cell *i* within the new coordinate system. Further details on the determination of the coordinate system and the mean shape are provided in Section 2 of S1 Appendix.

### Step 2: Vessel surface reconstruction

#### a) Estimation of vessel cross-sections

We estimated *M* local cross-sectional shapes *φ*_*k*_(*u*) := *φ*(*u*; *θ*_*k*_) with parameters *θ*_*k*_, *k* = 1, 2, …, *M*, at equidistant positions *z*_1_ *< z*_2_ *<* … *< z*_*M*_ on the anterior-posterior axis. To homogenize the density of estimated vessel cross-sections for different vessel segment lengths, we scaled the number *M* of estimated cross-sectional shapes with the length of the segment on the anterior-posterior axis:

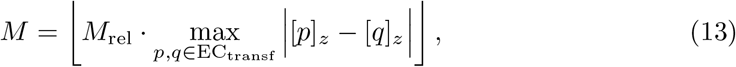

where *M*_rel_ is a tuning parameter controlling the relative number of cross-sections estimated per unit length (here: μm) for all data sets, i.e., embryos at either time point.

Each cross-sectional shape *φ*_*k*_ was initialized with the mean shape 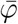. To account for the DA’s dorsal-ventral asymmetry, we employed superelliptic cross-sectional shapes. The superellipse shape interpolates between an ellipse and a rectangle (see Fig 3A) and is thus very suitable to model dorsal flattening. By joining two (half) superellipses, we allowed that the ventral segment of the DA’s cross-sections was not flattened. Our novel cross-sectional shape model

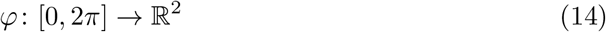

has parameters *θ* = (*m*_*x*_, *m*_*y*_, *a, b, c, α, β*) ∈ ℝ^7^ (see Fig 3B and also Section 3.1 of S1 Appendix).

**Fig 3.**
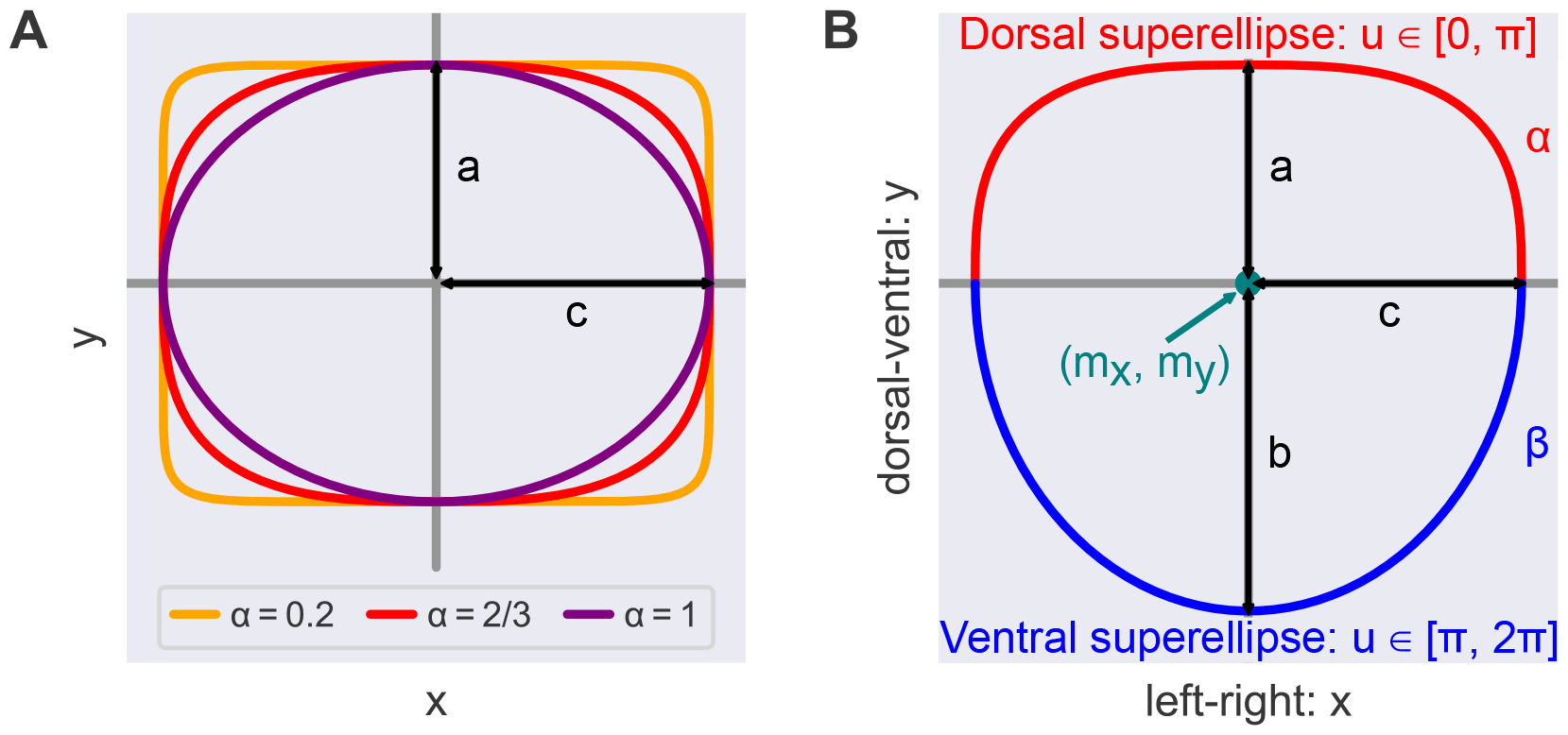
Cross-sectional shape model. **(A)** All points on the superellipses satisfy |*x/c*|^2*/α*^ + |*y/a*|^2*/α*^ = 1. For *α* = 1 the superellipse reduces to an ellipse. The smaller the value of *α <* 1, the closer the superellipse shape is to a rectangle. **(B)** We modeled DA cross-sections by joining two (half) superellipses. Here, (*m*_*x*_, *m*_*y*_) is the shape’s midpoint and *a, b, c* are the shape’s semi-axis lengths; *α, β* specify the extent of flattening of each superellipse.

A difficulty of the estimation of cross-sectional shapes is that there exists no closed form solution of the euclidean distance of a given point to a general superellipse. To obtain a simple and computationally inexpensive approximation of the orthogonal projection 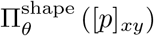 of the *xy*-coordinates of cell contour point *p* onto the cross-sectional shape *φ*(*u*; *θ*) with parameters *θ*, we approximated the shape by a sufficiently dense polygon consisting of *n*_poly_ points with linearly spaced parameter values (see also Section 3.2 of S1 Appendix).

We now explain the estimation of a local cross-sectional shape (see Fig 4 for an exemplary illustration). EC junction markers are only expressed on a small fraction of a blood vessel’s apico-luminal surface. Locally, information on cell junctions hence provides insufficient information to estimate any type of cross-sectional shape. To be able to estimate vessel cross-sectional shapes from cell junctions only, we considered all points on cell contour splines during each estimation: To evaluate the goodness of fit of the estimate *φ* with parameters *θ* on the cross-sectional plane at *z*_*k*_, we first computed the projections 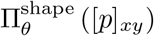 of the *xy*-coordinates of all cell contour spline points *p* ∈ EC_transf_ onto *φ*. We then defined the goodness of fit as a weighted sum of projection distances (see Eq (17)). To take into account that points closer to the cross-sectional plane carry more information on the local cross-sectional shape, we employed truncated Gaussian weights

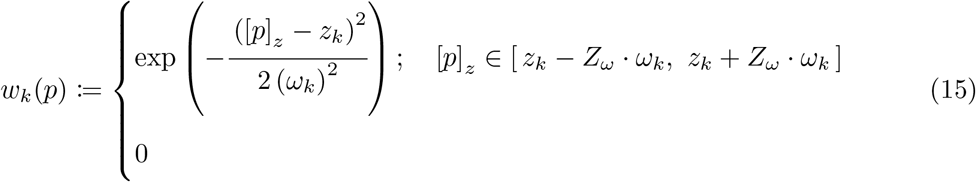

decaying with the distance of point *p* to the cross-sectional plane at *z*_*k*_. Here, *ω*_*k*_ *>* 0 and *Z*_*ω*_ *>* 0 are tuning parameters controlling the standard deviation and the distance of the truncation point in either direction from the mean value of the underlying Gaussian function, respectively.

**Fig 4.**
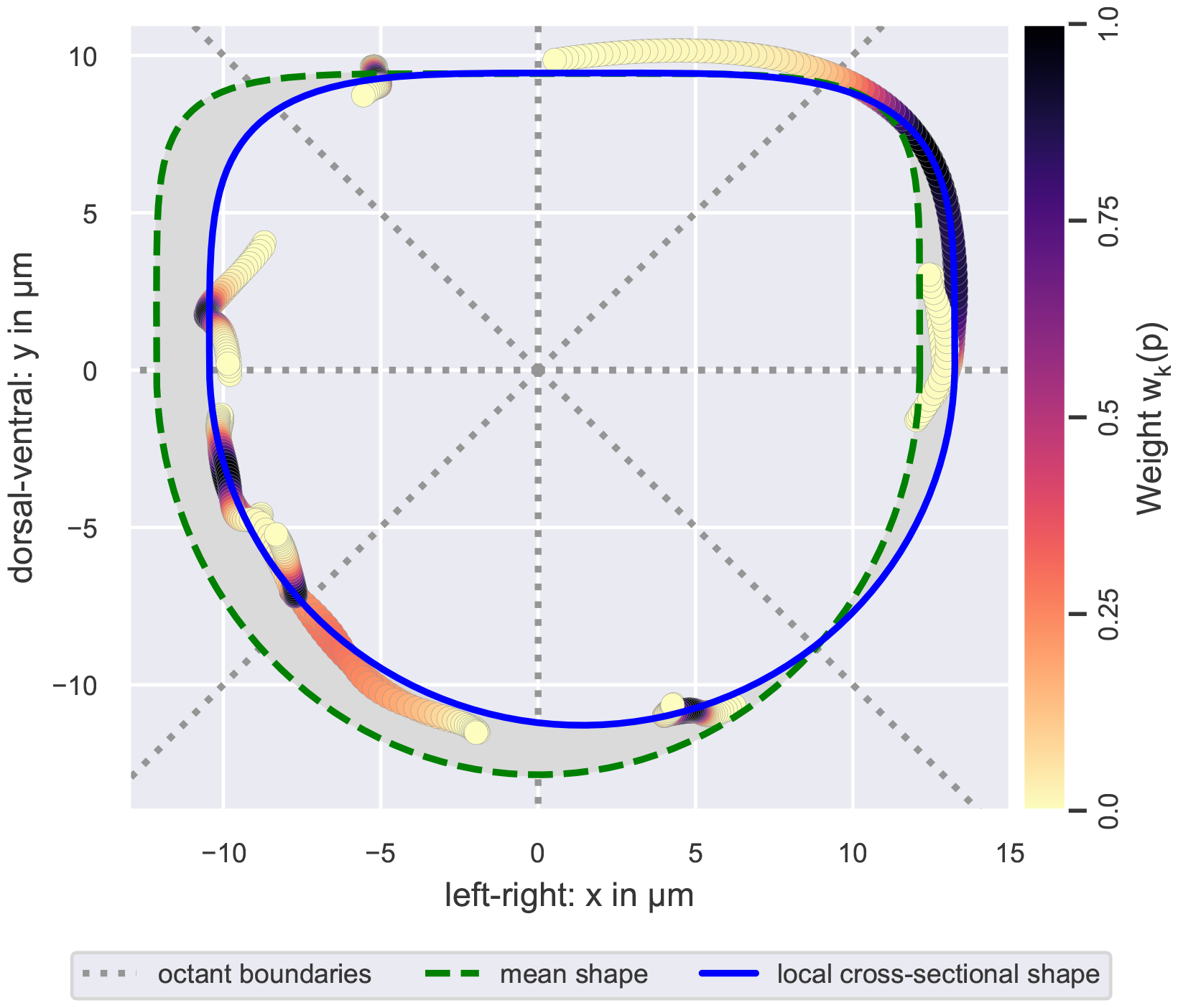
Estimation of a local cross-sectional shape. Visualized is the local cross-sectional shape estimated from points (circles) on EC contour splines that were projected onto the cross-sectional (*xy*-)plane. Data points are colored according to their weight during the estimation, i.e., their distance along the *z*-axis from the local cross-sectional plane (see color bar). Note that data points with a weight of zero are omitted in the figure. The employed weight function ensured that the estimation was based on at least *n*_oct_ data points with non-zero weight per plane octant. Initial parameter values for the local cross-sectional shape were provided by the mean shape. During estimation, the deviation of the local cross-sectional shape from the mean shape, i.e., the area of the gray region relative to the area within the green dashed curve, was bound from above by *λ*. The illustrated cross-sectional shape was estimated using *n*_oct_ = 30 and *λ* = 20 %.

The density of data points varied along the anterior-posterior axis, e.g., fewer data points were available at the annotation boundary than in the center of the anterior-posterior axis. Hence, we chose the width *ω*_*k*_ of the weight function in Eq (15) adaptive in space. To decide whether sufficient data points were available, we introduced the tuning parameter *n*_oct_ ∈ ℕ: We chose the minimal value for *ω*_*k*_ such that, when cell contour spline points EC_transf_ were projected onto the *xy*-plane, at least *n*_oct_ points *p* ∈ EC_transf_ with non-zero weight *w*_*k*_(*p*) were located in each octant of the cross-sectional plane. The plane octants are illustrated in Fig 4.

In some data sets, i.e., embryos at either time point, increasing the minimal number of points per plane octant is not sufficient to estimate a physiologically plausible cross-sectional shape: For example, if many points with high weights are located in ventral octants, but only points with low weights are located in dorsal octants, unconstrained estimation can result in cross-sectional shapes that fit closely to the ventral points but deviate strongly from the dorsal points (see for example Fig S6D). To avoid this local overfitting, we additionally constrained the deviation 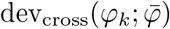 of the local cross-sectional shape *φ*_*k*_ from the mean shape 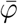 (see Eq (18)). We defined this deviation as the area of the symmetric difference of the two shapes, normalized by the mean shape’s luminal area:

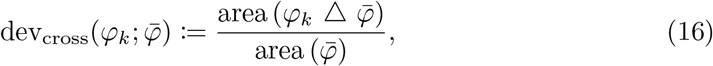

where △ is the symmetric difference of the interiors of two closed 2D curves (see gray region in Fig 4). To evaluate these symmetric differences, we approximated each curve by a polygon.

Eventually, we solved the following minimization problem for the estimation of the parameters *θ*_*k*_ of the cross-sectional shape at *z*_*k*_:

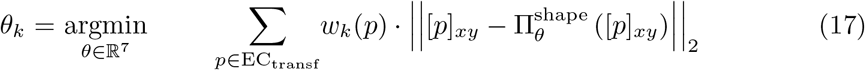

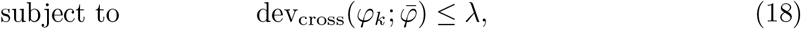

where *λ* ≥ 0 % is a tuning parameter that controls the strength of the constraint.

#### b) Smoothing of vessel surface

In Eq (17), the estimation of the different cross-sectional shapes is decoupled from each other; they only depend on the same input data. This drastically reduces the dimensionality of the optimization problem. Also, all optimization problems can be solved in parallel, which greatly reduces computation time. As a result, the transition of the estimated cross-sectional shapes, however, is not smooth along the anterior-posterior axis. To smoothen these transitions, we employed a Gaussian filter on the parameters of the cross-sectional shapes. This Gaussian filter is controlled by two tuning parameters: the standard deviation *σ >* 0 of the underlying Gaussian function and *Z*_*σ*_ *>* 0 that regulates the distance of the Gaussian kernel function’s truncation point in either direction from its mean value (comparable to *ω* and *Z*_*ω*_ in Eq (15), see also Section 4 of S1 Appendix). We denote the cross-sectional shape at *z*_*k*_ with smoothed parameters 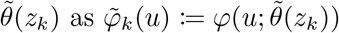.

#### c) Computation of grid

To later enable triangulation of cell surfaces, we approximated the reconstructed vessel surface as a grid (also called organized/ordered point cloud in the literature). To this end, we computed *n*_*k*_ equidistant points CS_*k*_ on each cross-sectional shape 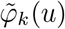 :

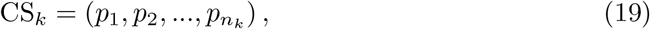

where points

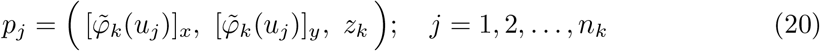

have parameter values

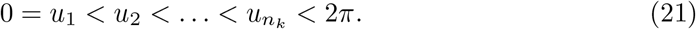

To homogenize the density of points computed on cross-sectional shapes with different circumferences, we scaled the number *n*_*k*_ of points computed per cross-sectional shape 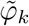 with its circumference:

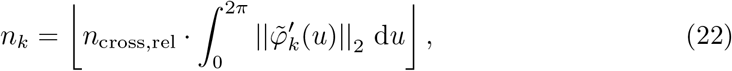

where *n*_cross,rel_ is a tuning parameter that controls the relative number of points computed per unit length (here: μm) for all cross-sections.

Analogous to the EC contours, we denote the successor/predecessor of a point *p* ∈ CS_*k*_ using *p*^+^ and *p*^−^, respectively:

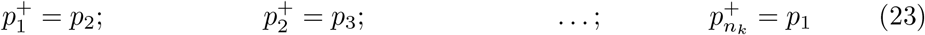

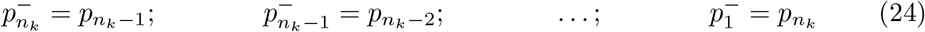

Throughout the text, it is typically clear from the context, whether *p*^+^ or *p*^−^ refers to neighbors within EC contours or within cross-sections. If this is not the case, clarifications are provided. Concatenating the points on all estimated cross-sections (*k* = 1, 2, …, *M*), an ordered sequence

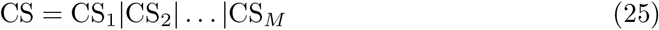

of length 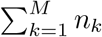 is obtained. We provide details on the computation of equidistant points on superelliptic shapes in Section 5.2 of S1 Appendix.

### Step 3: Endothelial cell surface triangulation

#### a) Projection of endothelial cell contours onto vessel surface

To define the EC surface boundaries on the estimated vessel surface, we projected each cell contour spline onto the grid approximation of the vessel surface. Whenever a spline *s*_transf,*i*_ intersected with a cross-sectional plane of the vessel at *z*_*k*_, the intersection point was determined. The projection of this intersection point onto the vessel surface was then approximated by the point’s nearest neighbor among the equidistant points CS_*k*_ on the local cross-sectional shape. We denote the points of the projected spline contour of cell *i* as EC_proj,*i*_. Details on the contour projection are provided in Section 6 of S1 Appendix.

#### b) Triangulation within projected endothelial cell contours

To obtain a smooth surface for each EC, we triangulated each cell individually ensuring that all edges of the projected cell contour were part of the triangulation. The triangulation was performed in-between each pair of neighboring cross-sections (see Fig 5): First, we collected all edges between subsequent points *p, q* ∈ EC_proj,*i*_ that connected cross-sections *k* and (*k* + 1), i.e., *p* ∈ CS_*k*_ and *q* ∈ CS_*k*+1_. If the edges connecting cross-sections *k* and (*k* + 1) were (*p*_1_, *q*_1_) and (*p*_2_, *q*_2_), we connected the cross-sectional paths within the cell surface between *p*_1_ and *p*_2_ and between *q*_1_ and *q*_2_ via triangles. Our custom triangulation algorithm is explained in detail in Section 7 of S1 Appendix.

**Fig 5.**
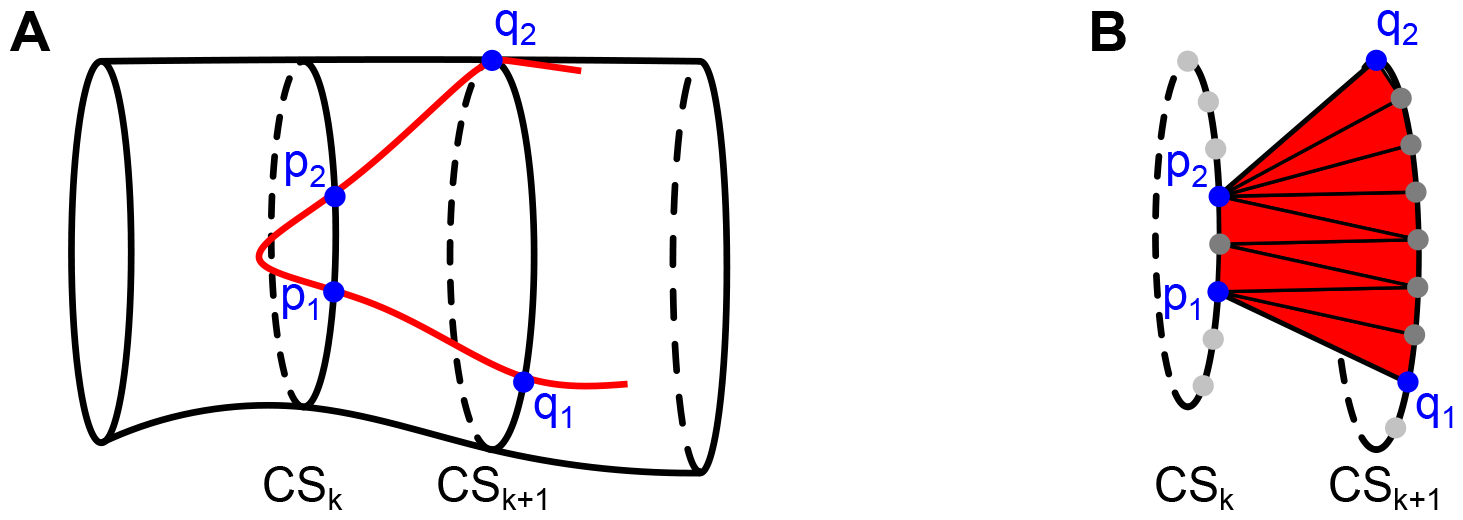
Triangulation within a projected cell contour. **(A)** In red, part of the projected cell contour on the smoothed vessel cross-sections is shown. To triangulate the part of the cell surface between cross-sections *k* and (*k* + 1), the edges (*p*_1_, *q*_1_) and (*p*_2_, *q*_2_) of the projected cell contour between these two cross-sections are identified. **(B)** A triangulation connects the cross-sectional path between *p*_1_ and *p*_2_ and the cross-sectional path between *q*_1_ and *q*_2_ within the cell surface. This triangulation is performed for each pair of neighboring vessel cross-sections that contains parts of the cell surface.

### Step 4: Analysis of vessel geometry and endothelial cell morphology

#### a) Quantification of cross-sectional geometry

We characterized cross-sectional geometry by two geometric measures:

1. The luminal area of the cross-sectional shape at *z*_*k*_ was approximated by the area of the polygon enclosed by cross-sectional points CS_*k*_:

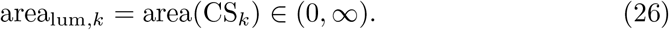
2. We defined the luminal diameter of the cross-sectional shape at *z*_*k*_ as the maximal distance between any two points *p, q* ∈ CS_*k*_ on the vessel cross-section:

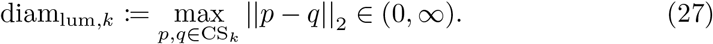

#### b) Quantification of endothelial cell surface morphology in 3D

We denote by *T*_*i*_ the set of triangles representing the surface mesh of endothelial cell *i*. Each triangle Δ ∈ *T*_*i*_ consists of three points in 3D space:

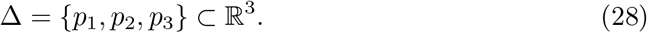

Based on this mesh, we characterized each endothelial cell surface by several morphometric measures:

1. The surface area of cell *i* was computed as the sum of the areas of its mesh’s triangles:

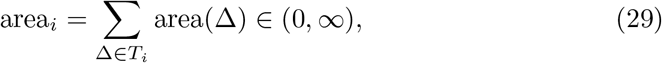

where area(Δ) is the area of triangle Δ.
2. The perimeter of cell *i* was computed as the sum of the lengths of the projected cell contour edges:

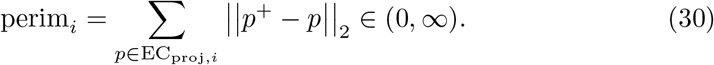
3. The compactness of cell *i* is defined as the ratio of the cell’s surface area to the area of a circle with the same perimeter:

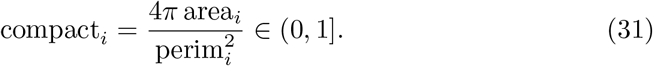 A circle has compactness 1, lower values correspond to elongated cells or cells with protrusions.
4. To quantify elongation of a cell surface in the direction of flow, we first defined a cell surface bounding box. This box was chosen such that it covered the cell’s extension within the cross-sectional plane and the cell’s length in the direction of flow. An example of this box is illustrated in Fig 6A.

We defined elongation as the ratio of the bounding box’s extension in the direction of flow (*z*) to its extension perpendicular to the direction of flow:

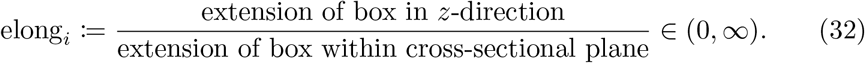

If elong_*i*_ *>* 1, cell *i* is elongated in the direction of flow; the higher the value, the more pronounced the elongation in the direction of flow. If elong_*i*_ ≤ 1, cell *i* is not elongated in the direction of flow; the lower the value, the more pronounced the elongation perpendicular to the direction of flow. We provide details on the computation of the bounding box and cell elongation in Section 8 of S1 Appendix.

**Fig 6.**
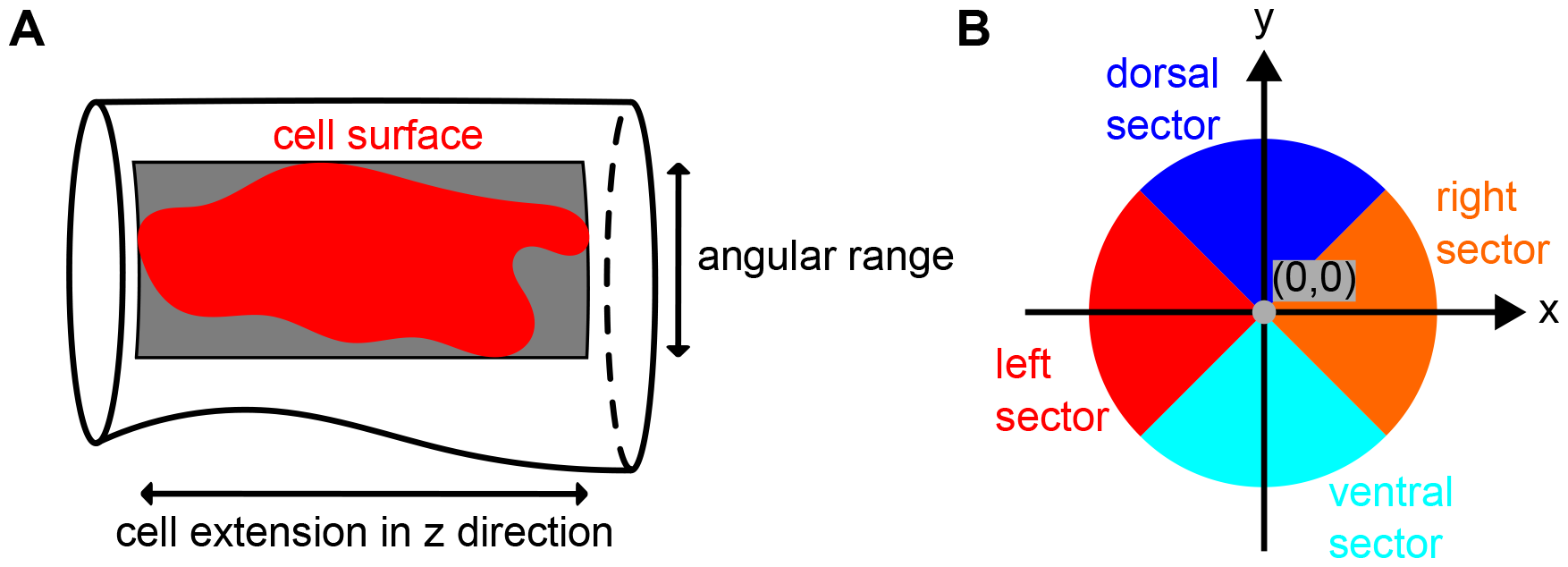
Details on characteristics used during morphological analysis. **(A)** The bounding box of a cell surface spans the entire angular range covered by the cell on the vessel surface along its length in *z*-direction. **(B)** Cross-sectional sectors for classification of cell location.

Finally, to quantify within-embryo variability in geometric or morphometric measures, we computed the quartile coefficient of dispersion (QCD):

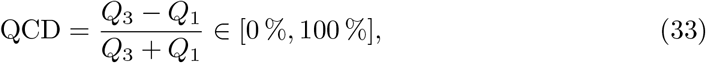

where *Q*_1_ and *Q*_3_ are the first and third quartiles of related measurements, e.g., EC surface areas, in an embryo at a single time point.

#### c) Classification of endothelial cells

We investigated whether ECs in different locations within the DA’s cross-sections differed in their morphological characteristics. For this, we divided the entire DA surface into dorsal, ventral, left and right sectors (see Fig 6B). If a cell’s surface area lying in the dorsal sector was greater than 50 % of its total surface area, we classified the cell as a dorsal cell; ventral cells were analogously classified. We did not distinguish between left and right cells. Thus, if a cell’s surface area lying in *either* the left *or* right sector was greater than 50 % of its total surface area, we classified the cell as a left/right cell. For details on the cell classification, see Section 9 of S1 Appendix.

### Goodness of fit

To study the impact of contour preprocessing and vessel surface reconstruction on the level of individual cell contours, we quantified the distances between the manually annotated EC contours EC_anno_ and their corresponding projected cell contours EC_proj_. To this end, we employed an outlier-insensitive and symmetric distance measure between pairs of contours.

Consider two contours 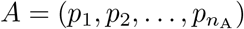 and 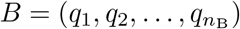 of points *p*_*j*_ ∈ ℝ^3^ with *j* = 1, 2, …, *n*_A_ and *q*_*j*_′ ∈ ℝ^3^ with *j*^′^ = 1, 2, …, *n*_B_. We first defined the distance *d*_*B*_(*p*) of a point *p* ∈ *A* to contour *B* as the minimal orthogonal distance to line segments (*q, q*^+^) with *q, q*^+^ ∈ *B* (see also Fig 7):

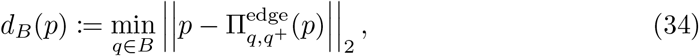

where 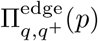 is the orthogonal projection of point *p* onto the line segment (*q, q*^+^) (see Eq (30) in S1 Appendix).

**Fig 7.**
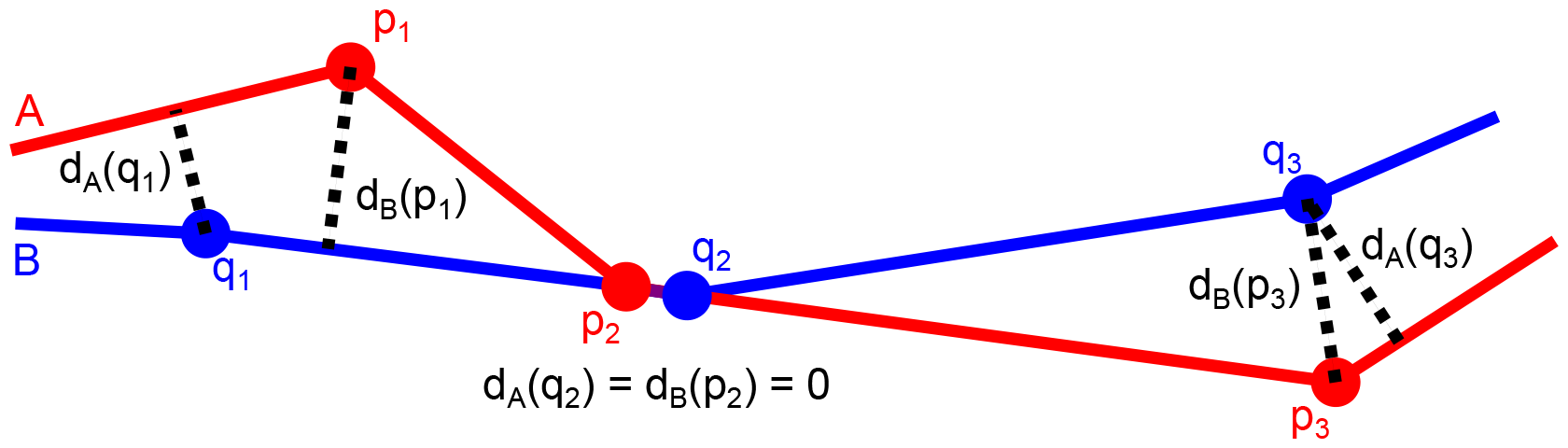
Point distances to contours. Distances (dotted lines) of points *p*_1_, *p*_2_ and *p*_3_ on contour A and points *q*_1_, *q*_2_ and *q*_3_ on contour B to the other contour (see also Eq (34)).

Next, we defined the asymmetric distance *d*_*B*_ (*A*) from contour *A* to contour *B* as the mean of the distances of points *p* ∈ *A* to contour *B*:

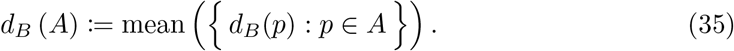

While using the median instead of the mean in Eq (35) would make the distance measure more robust to single point outliers, we found that employing the mean made the distance measure more robust when two contours resembled each other along most of their length but differed substantially along an entire segment of the contour. As the distances *d*_*B*_(*A*), *d*_*A*_(*B*) depend on the point densities of contours *A, B*, respectively, we linearly interpolated additional points on contours *A* and *B* prior to distance computations.

We finally defined the symmetric distance *d* (*A, B*) between contours *A* and *B* as the maximum of the two asymmetric distances *d*_*B*_ (*A*) and *d*_*A*_ (*B*):

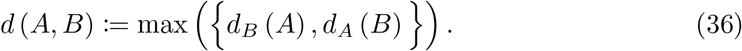

When the distance between a manually annotated EC contour and its projection onto the vessel surface is larger than the uncertainty stemming from the manual annotation process, this may indicate that (i) the chosen tuning parameter values for vessel surface reconstruction were not suitable (explained below), (ii) errors were introduced during manual annotation of the contour or (iii) the EC bends outwards from the DA as part of an intersegmental vessel.

### Criteria for choice of tuning parameters

During preprocessing of manually annotated EC contours and vessel surface reconstruction, we applied the *same* tuning parameter values to all wild-type and Endoglin-deficient embryos at both 48 hpf and 72 hpf. In short, tuning parameters controlling contour preprocessing were identified by visual inspection; each parameter’s magnitude depended only on the cell contours’ complexity, i.e., the cells’ deviation from an ellipse/hexagon due to irregular boundaries and protrusions: Suitable values for the parameters *r* and Δ*h*, which control EC contour enrichment by neighbors, ensure that the combined information of neighboring cells on a cell-cell contact is integrated into each of the cells’ contours. Values chosen for the parameters *n*_interp,rel_, *ϵ*_spline_ and *n*_spline,rel_, which control the approximation of EC contours by equidistant points on splines, should ensure that each EC contour is smoothed but still closely resembles its manual annotation. For further details, see Section 10.1 in S1 Appendix.

To precisely reconstruct local vessel geometry, we chose high values for the tuning parameters *M*_rel_, *n*_poly_ and *n*_cross,rel_ that control the number of estimated cross-sections along the vessel axis and the number of points computed on these cross-sections. Tuning parameters *Z*_*ω*_ and *Z*_*σ*_ controlling the truncation of the weight and smoothing functions were fixed to default values. The presented vessel surface reconstruction method balances the locality of estimated cross-sectional shapes, the physiological plausibility of these shapes and the overall smoothness of the vessel surface. These trade-offs are controlled by the three parameters *n*_oct_, *λ* and *σ*. For different combinations of the parameters’ values, we measured goodness of fit by computing the distances between the manually annotated EC contours EC_anno_ and their projections EC_proj_ onto the vessel surface using Eq (36). Additionally, we visually inspected the physiological plausibility of the estimated shapes and the smoothness of the resulting vessel surface. In large angular sectors of the cross-sectional plane that only contained points with low weights during the shape’s estimation, a physiologically plausible cross-sectional shape would either locally resemble the mean shape or closely match those points with low weights. For further details, see Section 10.2 in S1 Appendix.

To test whether the chosen values for the parameters *n*_oct_, *λ* and *σ* were robust, we performed a parameter sensitivity analysis. We investigated the effect of these parameters on vessel geometry and EC morphology by re-estimating each vessel surface with altered value of either *n*_oct_, *λ* or *σ*, while fixing the other two values. We then quantified the deviation dev_cross_(*φ*^∗^; *φ*) of each re-estimated cross-sectional shape *φ*^∗^ from its corresponding reference shape *φ* using Eq (16). Additionally, we quantified the relative deviation dev_rel_(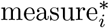 ; measure_*i*_) of each altered morphometric measurement 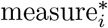, computed on a re-estimated vessel surface, from its corresponding reference value measure_*i*_ as

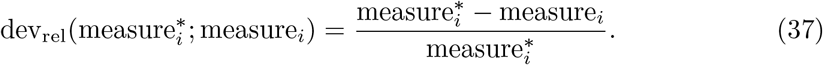

For example, if 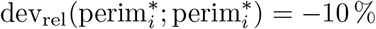, the EC perimeter 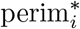, computed on a re-estimated vessel surface, is 10 % lower than its reference value perim_*i*_. We computed relative deviations of altered geometric measurements in the same manner.

### Model validation

To verify that our model enables accurate reconstruction of DA cross-sections using only information from EC contours, we compared the vessel surface estimated from EC contours in the DA of two wild-type embryos at 72 hpf to manually annotated angiogram slices of the same DAs (validation data in Table 1). During surface reconstruction, we used the same parameter values as in the primary analysis. We then compared the estimated cross-sectional geometry to the geometry of the annotated angiogram slices. As the planes containing annotated angiogram slices depended on the position of the embryos within the mounting medium during imaging and did not match cross-sectional planes in our estimated coordinate system, we sliced vessel surfaces estimated from cell contours with planes containing the angiogram slices. To then quantify differences in the (polygonal) shapes of vessel surface slice *S* and corresponding angiogram slice *S*_ref_, we computed their relative deviation dev_cross_(*S*; *S*_ref_) using Eq (16). Additionally, we quantified the relative deviation dev_rel_(area(*S*); area(*S*_ref_)) of their luminal areas using Eq (37).

Since prior methods had employed ellipses instead of superellipses to describe DA cross-sections [3, 15], we also investigated the impact of the employed cross-sectional shape model on the estimated vessel geometry. To this end, we re-estimated the vessel surfaces using either circular or elliptic cross-sectional shapes. To then quantify the deviation of the estimated circular or elliptic cross-sectional geometries from the geometry of the annotated angiogram slices, we again applied Eq (16) and Eq (37).

### Implementation and data availability

We implemented the reconstruction of zebrafish DA geometry and EC surface morphology from manually annotated EC contours in a python package called **CO**ntour-based **VE**ssel surface **R**econstruction (COVER). This package uses python 3.12, numpy [21], pandas [22] and scipy [23]. From scipy, we employed routines for minimization, root finding, quadrature, splines, nearest neighbor searches via KD trees and correlation analysis. We computed symmetric differences and polygonal areas with shapely [24]. Principal component analysis was performed using scikit-learn [25]. Furthermore, we employed pyvista [26] for visualization of triangular meshes, mesh clipping and mesh slicing. During method development and whenever inspecting intermediate results, we produced interactive plots with plotly [27]. All plots in the results section of this article, including its supplementary material, were created using matplotlib [28] and seaborn [29]. Our code, the manually annotated EC contours and the computed EC surface meshes are available from the Zenodo database via https://doi.org/10.5281/zenodo.10549102. We provide instructions to inspect the computed EC surface meshes using the open-source software Paraview [30] in S1 Tutorial.

## Results

Most results were computed using the analysis data (see Table 1); evaluations using the validation data are explicitly indicated.

### Robust choice of tuning parameter values

We chose tuning parameter values in accordance with the criteria summarized above. A summary of the parameters’ functions and their chosen values is provided in Table S2. Tuning parameter values used during contour preprocessing visibly reduced overlaps and gaps between neighboring cell contours, which we initially observed for the manually annotated cell contours. After vessel surface reconstruction, we computed the distances between the manually annotated EC contours EC_anno_ and their projections EC_proj_ onto the vessel surface using Eq (36). For the chosen tuning parameter values, we found that the resulting median contour distances were 45 % lower than those that were obtained when the vessel segment was instead described by a constant cross-sectional shape along its entire length (see Fig S6B–C). Additionally, we confirmed by visual inspection that the estimated vessel cross-sections had physiologically plausible shapes and together formed a smooth vessel surface (see exemplary illustrations in Fig S6D–E). Although we observed that tuning parameters *n*_oct_, *λ* and *σ* critically determined the shapes of estimated and smoothed vessel cross-sections (see exemplary illustrations in Fig S6D–E), chosen values for these parameters were robust: when halving or doubling either *n*_oct_, *λ* or *σ*, the majority of estimated vessel cross-sections had a relative deviation of less than 5 % (see Table S3) and no morphometric measurement differed by more than 10 % (see Table S4).

### Accurate projection of endothelial cell contours

For the chosen tuning parameter values, we analyzed whether the distances between the manually annotated cell contours EC_anno_ and their projections EC_proj_ onto the estimated vessel surface were sufficiently low to allow precise analysis of EC morphology. For this, we compared these distances with the uncertainty resulting from the manual annotation process: The distances between the manually annotated cell contours EC_anno_ (analysis data) and their projections EC_proj_ onto the estimated vessel surfaces had a median/maximum of 0.474 μm/1.36 μm, respectively. These distances were comparable to the median/maximal distances of 0.535 μm/0.742 μm found between the two annotation versions of the same cell contours in two wild-type embryos at 72 hpf (validation data; see Fig 8).

**Fig 8.**
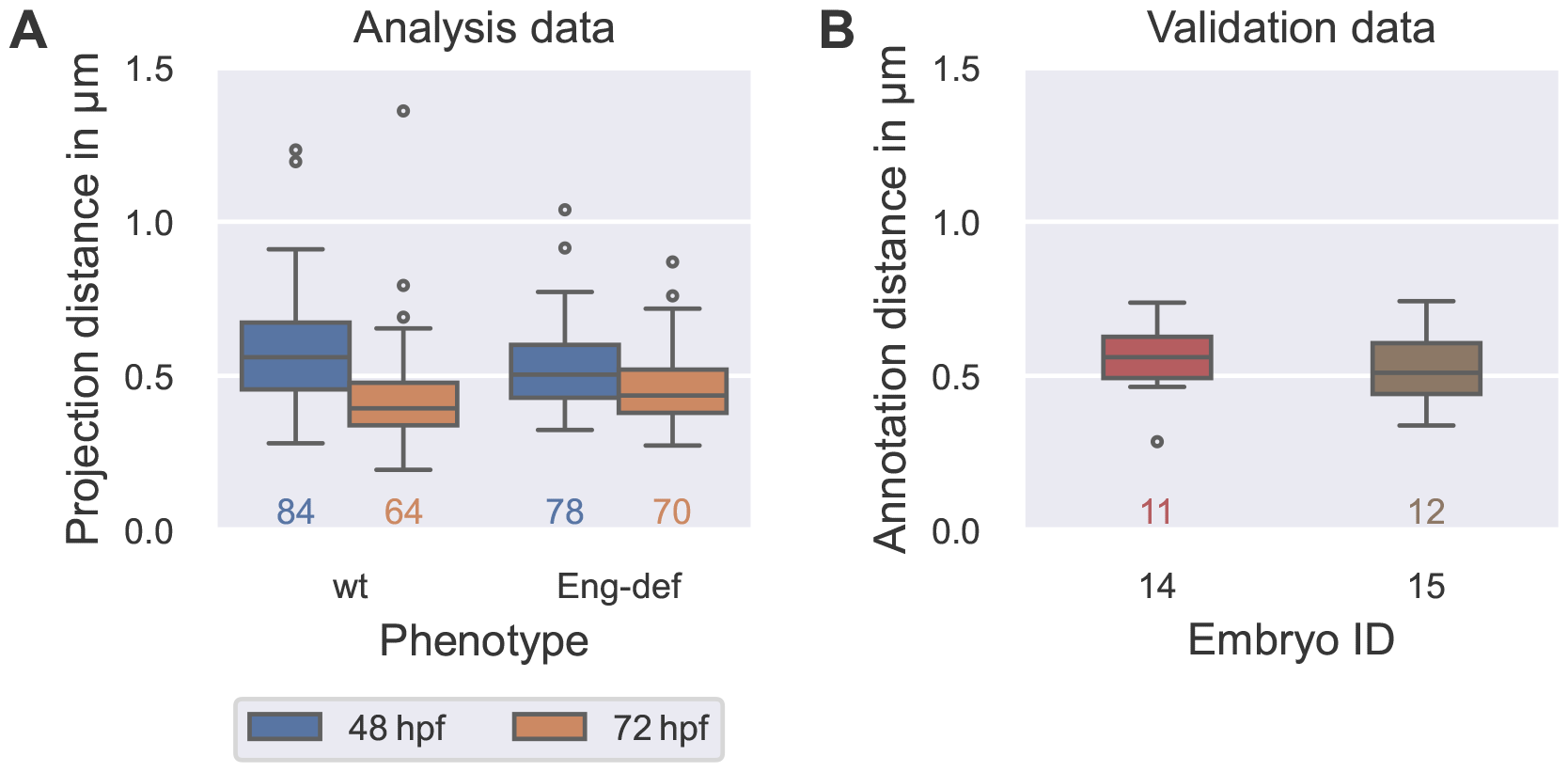
Comparability of errors introduced by vessel surface reconstruction and annotation uncertainty. **(A)** Distribution of the distances between manually annotated endothelial cell contours EC_anno_ (analysis data) and their projections EC_proj_ onto the estimated vessel surface (see Eq (36)), stratified by phenotype and time. **(B)** Distribution of distances (see Eq (36)) between first and second annotation of endothelial cells in two wild-type embryos at 72 hpf (validation data). The numbers below each box plot denote the corresponding numbers of cells.

We inspected ECs with high projection distances (analysis data). Here, we found that these were cells that partially bend outwards from the DA as part of an intersegmental vessel or cells at the annotation boundary where, locally, an insufficient number of data points was available to accurately estimate local cross-sectional shapes. To ensure that we only quantified morphology in ECs whose contours were well represented on the estimated vessel surface, we excluded those 22 out of 296 cells that had projection distances greater than the maximal annotation uncertainty of 0.742 μm in the following morphological analysis (from the analysis data).

### Accurate estimation of vessel cross-sections from cell contours

After confirming that our method was precise enough for the computation of EC morphology, we evaluated whether the estimation of vessel cross-sections using only information from EC contours resulted in an accurate description of DA geometry. For this, we compared vessel surfaces estimated from cell contours of two wild-type embryos at 72 hpf to angiogram slices from the same DAs (validation data). Pooling both embryos, we found that 92 % to 93 % of vessel surface slices had luminal areas differing by at most 20 % from the areas of angiogram slices when using either circular, elliptic or superelliptic cross-sections. For a more detailed analysis of luminal area in each of the two embryos, see Table S5.

We further found that the *shapes* of estimated vessel cross-sections resembled the shapes of angiogram slices well for each tested cross-sectional shape. Ellipses and superellipses, however, were slightly more accurate in describing angiogram slices than circles: when pooling both embryos, the median deviation of vessel surface slices from the angiogram slices was 17 %/14 %/15 % when using circular/elliptic/superelliptic cross-sections, respectively. For a more detailed analysis of luminal shape in each of the two embryos, see Fig S7A.

Using the same data, we then investigated how the density of annotated cell contours affected the approximation quality of the estimated (superelliptic) vessel cross-sections from the angiogram slices. Pooling both embryos, the median deviation of the estimated cross-sections from the angiogram slices was 61 % higher for angiogram slices where only one annotated EC was located within a distance of 0.5 μm in comparison to slices with more nearby cells. For a more detailed analysis of the effect of annotation density on the deviation in luminal shape in the individual embryos, see Fig S7B. The effect of annotation density on the accuracy of cross-sectional shapes motivated us to exclude all cross-sections which had only one annotated cell within a distance of 0.5 μm to the cross-sectional plane (3697 out of 25 111 cross-sections) in the subsequent analysis of DA geometry (from the analysis data).

### High between-embryo but low within-embryo variability in luminal area

Once we confirmed that our method accurately estimated vessel cross-sections using only information from EC contours, we quantified cross-sectional geometry in the DA of wild-type and Endoglin-deficient embryos at 48 hpf and 72 hpf (see Fig 9). We found that median luminal area in the pooled Endoglin-deficient embryos was 34 % smaller at 48 hpf in comparison to the pooled wild-types, but 150 % larger at 72 hpf. Comparing the individual embryos, we found that biological between-embryo variability in luminal area was remarkably high in both phenotypes at each time point: the maximal median luminal area over all wild-type embryos was 90 %/49 % higher than the corresponding minimal median luminal area at 48 hpf/72 hpf, respectively (compare Fig 9A). Among Endoglin-deficient embryos, we found that the maximal median luminal area was 71 % higher than the minimum at 48 hpf and 57 % higher at 72 hpf (compare Fig 9B). In contrast to the high between-embryo variability, biological *within*-embryo variability in luminal area was low in the analyzed short segments of the DA: the maximal QCD (see Eq (33)) over all embryos and time points was only 11 %.

**Fig 9.**
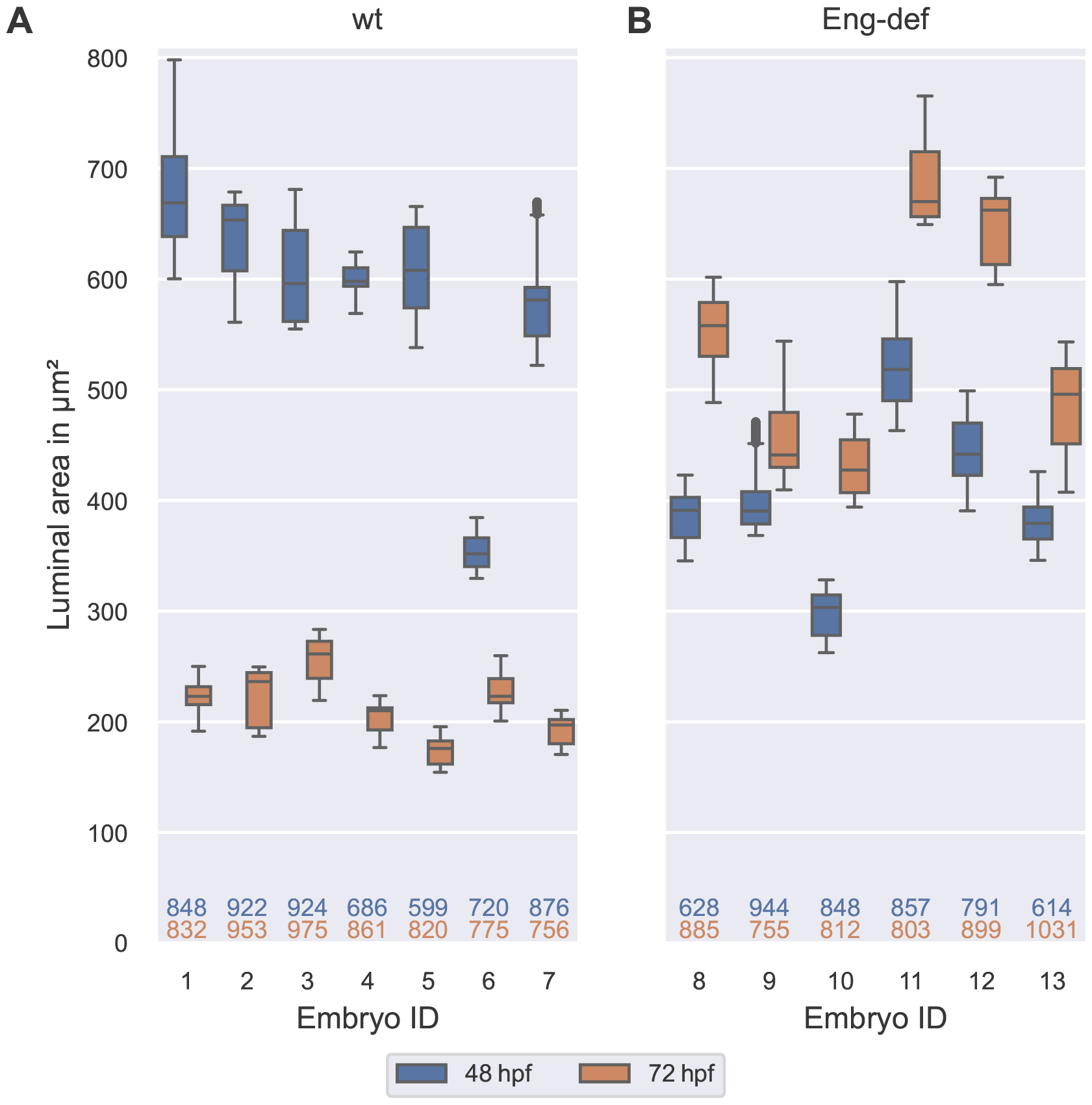
High between-embryo but low within-embryo variability in luminal area. Distribution of luminal areas of estimated DA cross-sections, stratified by embryo and time, for wild-type embryos **(A)** and for Endoglin-deficient embryos **(B)**. Analysis based on cross-sections with at least two annotated cells present within a distance of 0.5 μm to the cross-sectional plane. The numbers below the box plots denote the corresponding numbers of vessel cross-sections. The top number corresponds to 48 hpf and the bottom number to 72 hpf.

When inspecting each embryo individually, we found that in each wild-type embryo, median luminal area decreased between 48 hpf and 72 hpf (see Fig 9A), while it *increased* in each Endoglin-deficient embryo (see Fig 9B). Relative changes in median luminal area between 48 hpf and 72 hpf however varied per embryo. For example, wild-type embryo 6, which already had the lowest median luminal area at 48 hpf, had a much smaller relative decrease in median luminal area of 37 % in comparison to the other wild-type embryos where the relative decreases ranged from 56 % to 71 %. Changes in luminal area also varied between Endoglin-deficient embryos. Here, increases in median luminal area ranged from 13 % to 50 %. In summary, we found high between-embryo but *low within*-embryo variability in luminal area in short DA segments of both wild-type and Endoglin-deficient embryos.

### High between-embryo and high within-embryo variability in endothelial cell morphology

Following our evaluation of cross-sectional geometry, we quantified EC morphology in the same DAs of wild-type and Endoglin-deficient embryos at 48 hpf and 72 hpf (see Fig 10). We found that median EC surface area in the pooled Endoglin-deficient embryos was 21 % smaller at 48 hpf in comparison to the pooled wild-types, but 40 % *larger* at 72 hpf. Inspecting changes in EC surface area between 48 hpf and 72 hpf, we observed opposing trends in the two phenotypes: Among the wild-type embryos, embryos 1 and 3 were the only embryos where median cell surface area increased (by 43 % and 6.6 %, respectively; see Fig 10A). In the remaining five wild-types, *decreases* in median cell surface area ranged from 3.7 % to 34 %. In contrast, median cell surface area substantially *increased* in almost all Endoglin-deficient embryos; these increases ranged from 43 % to 170 % (see Fig 10B). Embryo 13 was the only Endoglin-deficient embryo where median cell surface area slightly *decreased* between 48 hpf and 72 hpf (by 17 %).

**Fig 10.**
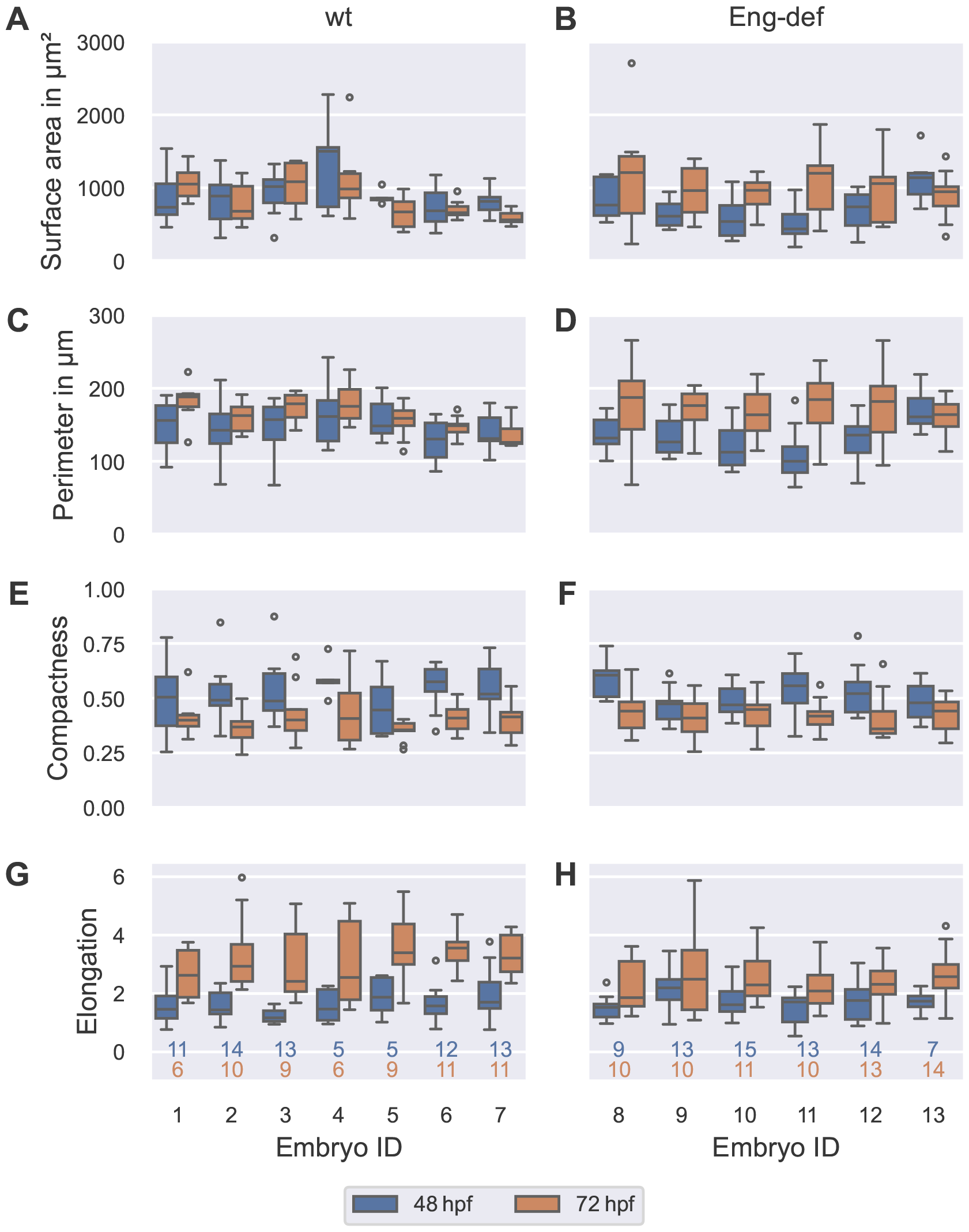
High between-embryo and high within-embryo variability in endothelial cell morphology. Distribution of morphometric measurements over all ECs, stratified by embryo and time, for wild-type embryos (left column) and for Endoglin-deficient embryos (right column). Analysis based on ECs with projection distances (onto the estimated vessel surfaces) comparable to or lower than the maximal annotation uncertainty of 0.742 μm. The numbers below the bottom box plots denote the corresponding numbers of cells for each column. The top number corresponds to 48 hpf and the bottom number to 72 hpf.

Although we observed clear differences in EC surface area between the two phenotypes, perimeters were comparable! Median EC perimeter in the pooled Endoglin-deficient embryos differed by maximally 11 % from that of the pooled wild-type embryos at each time point. When inspecting the individual embryos, we found that median EC perimeters increased in almost all embryos between 48 hpf and 72 hpf; the only *decrease* (by 4.0 %) occurred in wild-type embryo 7 (see Fig 10C–D). Notably, increases in median perimeter were greater in Endoglin-deficient embryos: Excluding embryo 13 where median perimeter only increased by 1.9 %, median perimeter increases ranged from 34 % to 85 % in the remaining Endoglin-deficient embryos, but only from 7.0 % to 21 % in wild-types.

Compactness of ECs was also similar in the two phenotypes: median compactness was 2.4 % smaller in the pooled Endoglin-deficient embryos than in the pooled wild-types at 48 hpf but 7.4 % larger at 72 hpf. When comparing individual changes in compactness between 48 hpf and 72 hpf, we found that median compactness decreased in each embryo (see Fig 10E–F). Between-embryo variability in this decrease was however higher in Endoglin-deficient embryos: Here, decreases in median compactness ranged from 4.5 % to 31 %, while decreases in wild-type embryos only ranged from 18 % to 30 %.

In comparison to the pooled wild-types, median EC elongation was 19 % higher in the pooled set of Endoglin-deficient embryos at 48 hpf, but 28 % *lower* at 72 hpf. Notably, median elongation increased in each embryo between 48 hpf and 72 hpf (see Fig 10G–H). These increases were however less pronounced in Endoglin-deficient embryos where they ranged from 14 % to 48 %, while increases in wild-type embryos ranged from 74 % to 130 %.

To analyze whether there were systematic relationships between any of the computed morphometric measurements, we next performed a correlation analysis (see Fig S8). When pooling all ECs, we found that cell surface area and perimeter showed a strong positive correlation (Spearman rank correlation coefficient *r*_*s*_ = 0.849). Also, when separating cells by phenotype and time, this correlation remained strong (0.784 ≤ *r*_*s*_ ≤ 0.916). This meant that ECs had few protrusions, otherwise large increases of perimeter could have been accompanied by small increases in surface area. Pooling all ECs, elongation and compactness had a moderately strong negative correlation (*r*_*s*_ = −0.640). When separating cells by phenotype and time, this correlation was slightly weaker (−0.562 ≤ *r*_*s*_ ≤ −0.466). In the absence of many protrusions, this negative correlation implied that very elongated cells (with low compactness) had a tendency to be aligned in the direction of flow, i.e., they had a high value of our elongation measure.

Finally, we analyzed within-embryo variability in morphometric measures. Notably, we found that biological within-embryo variability in EC morphology was much higher than within-embryo variability in luminal area in the same DA (see Fig S9). This was most evident in EC surface area and elongation: In 96 %/100 % of data sets, i.e., embryos at either time point, variability (QCD) in EC surface area/elongation was more than 50 % higher than QCD of luminal area, respectively. These two morphometric measures also had systematically higher within-embryo variability than EC perimeter and compactness (see Fig S10): In 85 %/65 % of data sets, QCD of EC surface area was more than 25 % higher than QCD of perimeter/compactness, respectively. In 77 % of data sets each, QCD of EC elongation was more than 25 % higher than QCD of perimeter and compactness. In summary, we found high between-embryo and high within-embryo variability in EC morphology in both wild-type and Endoglin-deficient embryos. Among the morphometric measures, EC surface area and elongation in the direction of flow showed the greatest within-embryo variability.

### Dorsal-ventral asymmetry of endothelial cell morphology

We then analyzed whether stratification of cells by their location within the DA reduced the remaining high within-embryo variability in EC morphology, especially in EC surface area and elongation. For 253 out of 274 ECs, we were able to identify the cross-sectional sectors containing the largest part of the cells’ surface area. This allowed us to study morphological differences between dorsal, ventral and left/right ECs in both wild-type and Endoglin-deficient embryos (see Fig 11).

**Fig 11.**
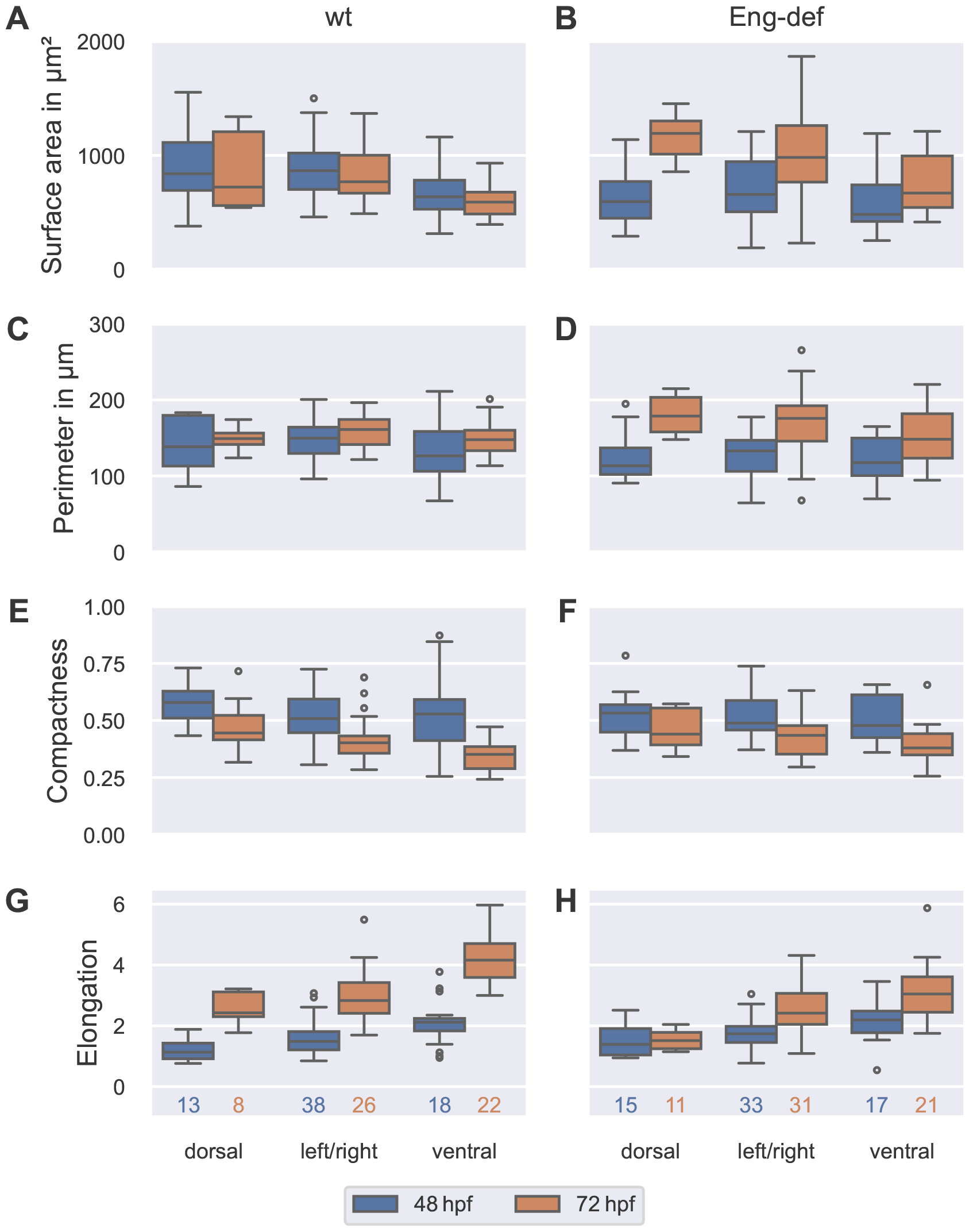
Dorsal-ventral asymmetry of endothelial cell morphology. Distribution of morphometric measurements over all ECs, stratified by location and time for wild-type embryos (left column) and for Endoglin-deficient embryos (right column). Analysis based on ECs with projection distances (onto the estimated vessel surfaces) comparable to or lower than the maximal annotation uncertainty of 0.742 μm. The numbers below the bottom box plots denote the corresponding numbers of cells for each column.

In wild-type embryos, we found evidence of a dorsal-ventral asymmetry. Firstly, dorsal ECs were larger than ventral ECs: Median surface area of dorsal ECs was 32 %/22 % larger at 48 hpf/72 hpf, respectively (see Fig 11A). Differences in perimeter were less pronounced; median perimeter of dorsal ECs was 9.5 % higher at 48 hpf but only 1.0 % higher at 72 hpf (see Fig 11C). We further observed that dorsal ECs were more compact than ventral ECs but *less* elongated in the direction of flow: Median compactness of dorsal ECs was 9.6 % larger at 48 hpf and 27 % larger at 72 hpf (see Fig 11E). In contrast, median elongation was 47 %/42 % *lower* in dorsal ECs at 48 hpf/72 hpf, respectively (see Fig 11G). When investigating changes in these measures over time, we found that dorsal and ventral ECs differed most in surface area and perimeter: Between 48 hpf and 72 hpf, median surface area of dorsal ECs decreased by 14 %, but ventral ECs only showed a decrease of 7.3 % (see Fig 11A). While median perimeter only increased by 7.8 % in dorsal ECs, its increase was 17 % in ventral ECs (see Fig 11C).

Our analysis of EC morphology in Endoglin-deficient embryos also supported a dorsal-ventral asymmetry. We found that dorsal ECs were larger than ventral ECs at both time points, but this difference was much more pronounced at 72 hpf: In comparison to ventral ECs, median cell surface area of dorsal ECs was 23 % larger at 48 hpf but 78 % larger at 72 hpf (see Fig 11B). While median perimeter of dorsal ECs was 3.4 % smaller at 48 hpf, it was 21 % *larger* at 72 hpf (see Fig 11D). Again, we observed that dorsal ECs were more compact than ventral ECs but *less* elongated in the direction of flow: Median compactness of dorsal ECs was 11 % larger at 48 hpf and 16 % larger at 72 hpf (see Fig 11F). In contrast, median elongation was 37 %/50 % *lower* in dorsal ECs at 48 hpf/72 hpf, respectively (see Fig 11H). Dorsal and ventral ECs not only had large differences in cell surface area and elongation at both 48 hpf and 72 hpf but changes in these measures over time also differed in their magnitude: Between 48 hpf and 72 hpf, median surface area increased by 100 % in dorsal ECs but only by 39 % in ventral ECs (see Fig 11B). Simultaneously, median elongation of dorsal ECs barely increased (by 9.0 %), while median elongation of ventral ECs increased by 39 % (see Fig 11H).

As differences between wild-type and Endoglin-deficient embryos were most pronounced in EC surface area and elongation, we finally compared surface area and elongation in dorsal and in ventral ECs between the two phenotypes. For each of the two morphometric measures, we found that dorsal ECs showed the greatest differences between the two phenotypes: At 48 hpf, median cell surface area of dorsal ECs was 29 % smaller in Endoglin-deficient embryos than in wild-types, while median cell surface area of ventral ECs was only 24 % smaller in Endoglin-deficient embryos (see Fig 11A–B). At 72 hpf, differences between the two phenotypes were even more pronounced in dorsal ECs: here, median cell surface area of dorsal ECs was 65 % larger in Endoglin-deficient embryos than in wild-types, while median cell surface area of ventral ECs was only 13 % larger in Endoglin-deficient embryos. Comparing elongation between the two phenotypes, we found that median elongation of dorsal ECs was 22 % higher in Endoglin-deficient embryos at 48 hpf (see Fig 11G–H). In contrast, median elongation of ventral ECs of Endoglin-deficient embryos was only 3.5 % higher. At 72 hpf, both dorsal and ventral ECs showed greater differences between the two phenotypes: here, median elongation of dorsal ECs was 38 % *lower* in Endoglin-deficient embryos than in wild-types, while median elongation of ventral ECs was only 27 % lower.

Finally, we closely inspected those ECs that could not be classified (see Fig S11). These cells contained cell surface areas in at least three of the dorsal, ventral, left and right sectors. Most of the cells had the largest surface area contribution in the dorsal sector (14 out of 21 ECs); no EC had the largest surface area contribution in the ventral sector. In each phenotype and at each time point, non-classifiable ECs had higher median cell sizes than dorsal, ventral and left/right ECs. Median compactness hardly differed between non-classifiable ECs and the classified ECs. Lastly, non-classifiable ECs had the lowest median elongation in the direction of flow. Taken together, the large size of the non-classifiable ECs, their low elongation in the direction of flow and predominant location in the dorsal sector further support a dorsal-ventral asymmetry of endothelial cell morphology. In summary, we found that dorsal and ventral ECs in the DA of both wild-type and Endoglin-deficient embryos vastly differed in their morphology, especially in their surface areas and elongation in the direction of flow. Furthermore, dorsal ECs showed the most pronounced differences between wild-types and Endoglin-deficient embryos.

## Discussion

In this article, we developed a novel mathematical morphology model of the zebrafish DA that enables reconstruction of the vessel’s apico-luminal surface using only data from annotated EC contours and approximate directions of the vessel’s anterior-posterior, dorsal-ventral and left-right axes; information from EC surface markers is not required. We found that errors on the EC contour level that were introduced by vessel surface reconstruction were comparable to the uncertainty stemming from the manual contour annotation process. Furthermore, we validated the accuracy of our EC contour-based vessel surface reconstruction method by comparing the luminal shapes of estimated vessel surfaces of wild-type embryos against angiography of the same DAs. Application of our model enabled us to precisely and robustly quantify DA geometry and 3D EC surface morphology in wild-type and Endoglin-deficient zebrafish embryos at 48 hpf and 72 hpf.

To ensure a physiologically accurate description of the DA’s surface, we developed a novel cross-sectional shape model that explicitly accounts for the vessel’s dorsal flattening that we had observed in 3D microscopy images. We found that vessel surfaces estimated using the novel superelliptic shape model described angiography data of the DA similarly well as frequently employed ellipses (see Fig S7A). Compared with ellipses, however, our shape’s dorsal-ventral asymmetry can not only accurately describe the DA’s geometry but also aid in identification of a physiologically plausible coordinate system of the vessel from initial direction vectors, while neither the rotation angle nor the semi-axis lengths of an ellipse are informative with respect to the vessel’s dorsal-ventral axis. Independent of the employed cross-sectional shape, an alternative way to identify the DA’s dorsal-ventral axis is to leverage further spatial information, e.g., the positions of ECs of the ventrally located caudal vein as done in [15]. While both approaches result in physiologically meaningful coordinate systems that allow to compare DAs in different embryos, our method does not require any data from neighboring vessels.

A major challenge during vessel surface reconstruction was that information on the left and right side of the DA was imbalanced: Since embryos lay on their right side within the confocal microscope, their left side was further away from the light source and therefore prone to bleaching effects and signal scattering, resulting in a loss of resolution and fluorescence signal; this made EC contour annotation on the left more difficult. This was more prominent for embryos at 48 hpf, since their larger yolk sacks increased the distances of their DAs from the microscope. Here, our novel cross-sectional shape model again aided in the estimation process: Whenever cell contours on the DA’s left side could not be annotated with confidence (and were thus omitted from the analysis), the shape’s symmetry axis and employing constrained minimization helped to estimate physiological cross-sectional shapes from an incomplete set of local cell contours. Freely rotating ellipses cannot accomplish this.

By first estimating vessel cross-sections from EC contours and then projecting each contour onto the estimated cross-sectional shapes, our model was not only able to describe variability in EC surface morphology but also explain it by the cells’ location within the vessel cross-sectional plane. While the former is also possible when only estimating individual EC surfaces like in [3], the latter additionally requires the reconstruction of vessel geometry in a manner that is consistent with all ECs. Since we based the estimation of vessel cross-sections on the combined information of *all* EC contours, our model can additionally compensate for annotation errors in single cell contours by contours of cells that belong to the same vessel cross-sections. The EC surfaces produced by our method are thus less sensitive to uncertainty stemming from the manual annotation process than EC surfaces whose estimation is only informed by single cell contours like in [3].

When comparing measurements made on the reconstructed surfaces, we found that the quantified vessel geometry agreed well with the literature [3, 16] (see Table S6): Mean luminal areas of the DA in wild-type zebrafish embryos at 48 hpf and 72 hpf differed less than 5 % from the mean areas reported in [16]. When additionally computing DA diameters with Eq (27), we found that mean DA diameters of wild-type and Endoglin-deficient embryos at 48 hpf and 72 hpf maximally differed by 14 % from the mean diameters measured in [3].

Comparing EC morphology, however, we found that our computed mean cell surface areas and perimeters were consistently smaller in each phenotype and at each time point than those in [3] (see Table S7). At 72 hpf, these differences were more pronounced; most notably, mean surface areas of wild-type embryos were 24 % smaller than those reported in [3]. While we observed the same trend of increasing EC surface area and perimeter of Endoglin-deficient embryos between 48 hpf and 72 hpf, the authors in [3] measured a higher relative increase (of 71 %) in mean EC surface area of these embryos between 48 hpf and 72 hpf than we did (48 %). Their consistently greater EC sizes could potentially be explained by differences between the developmental stages of the embryos analyzed in this article and those in [3]: In any study, the process of imaging an entire group of zebrafish embryos takes several hours. These delays during image acquisition increase between-embryo variability. Alternatively, the observed differences could be due to differences in data acquisition, e.g., due to differences in the experimental setup or uncertainty during manual annotation. To investigate whether our surface reconstruction could have impacted EC morphometrics, we additionally computed perimeters from the manually annotated EC contours prior to contour preprocessing and surface reconstruction. The perimeters of the EC contours that were projected onto the estimated vessel surface had a maximal deviation of 12 % from the perimeters of the corresponding manually annotated EC contours, respectively. Thus, the observed differences in cell perimeter were not introduced by our vessel surface estimation

Although the authors in [16] did not incorporate information on EC contours in their model, they were able to compute averaged EC surface areas from the DA’s surface area and the number of EC nuclei within the DA. While their mean EC surface area fit the mean value of our measurements well at 72 hpf (relative deviation of 6.2 %), they measured a lower mean EC surface area at 48 hpf (relative deviation of 42 %; see Table S7): Our larger mean EC size at 48 hpf could be explained by our decision to exclude (small) ventral cells that were extruding from the DA during annotation. As these cells are leaving the DA, they no longer contribute to vessel geometry.

In our analysis we quantified a previously unrecognized dorsal-ventral asymmetry of EC morphology in both wild-type and Endoglin-deficient DAs. Two distinct mechanisms could explain this dorsal-ventral asymmetry: The cell morphology of dorsal and ventral cells could be regulated or constrained by their respective biomechanical environment, e.g., by the presence of the dorsally located stiff notochord or the ventrally located compliant caudal vein (see the mechanical model in [16]). Alternatively, it is possible that dorsal and ventral ECs experience different levels of wall shear stress (WSS). In [31], it was shown that endothelial overexpression of the flow-sensitive transcription factor Klf2a decreased endocardial cell size in zebrafish embryos. Hence, if ventral ECs in the DA experienced higher levels of WSS, this could explain their smaller cell size. This hypothesis is further supported by our finding that, in both phenotypes, ventral ECs were more strongly elongated in the direction of blood flow than dorsal ECs, a process that is mediated by WSS [32]. It is up to follow-up live imaging studies to investigate whether there exists a correlation or even a causation between the observed dorsal-ventral asymmetry of EC morphology in the DA and local levels of wall shear stress in wild-type and Endoglin-deficient embryos. Functional studies could further explore whether the observed morphological differences between dorsal and ventral ECs in Endoglin-deficient embryos are directly linked to distinct local activity levels of Endoglin and, if that is the case, how the protein’s activity is affected by local wall shear stress. Insights generated from such studies can aid in finding out whether variability in EC shape also contributes to the onset of human arteriovenous malformations that occur due to a loss of Endoglin [12].

In summary, we developed a novel mathematical morphology model of the zebrafish DA that allows to reconstruct the vessel’s apico-luminal surface using only information from EC contours. Our model’s consistent description of vessel geometry and EC morphology enables it to generate new insights into spatial variability in EC morphology and the relationship between EC morphology and vessel geometry. We demonstrated this ability by applying our model to wild-type and Endoglin-deficient zebrafish embryos where we identified a previously unrecognized dorsal-ventral asymmetry of EC morphology. Notably, we found that dorsal ECs contributed most to the vessel diameter increase that occurs in Endoglin-deficient embryos between 48 hpf and 72 hpf.

## Supporting information

**S1 Appendix Supporting methods**. The text includes Fig S1 – Fig S5 and Table S1.

**S2 Appendix Supporting results**. The file bundles Fig S6 – Fig S11 and Table S2 – Table S7.

**S1 Tutorial Inspecting endothelial cell surface meshes using Paraview**.

**Fig S1 Insertion of points from neighboring cell contours during enrichment.**

**Fig S2 Coordinate system and mean shape.**

**Fig S3 Projection of cell contour spline onto vessel surface. Fig S4 Triangulation respecting cell contour edges.**

**Fig S5 Extraction of triangulation within cell contour. Fig S6 Iterative choice of tuning parameter values.**

**Fig S7 Compliance of estimated cross-sectional shapes with angiography. Fig S8 Correlations between morphometric measures.**

**Fig S9 Within-embryo variability in endothelial cell morphology larger than within-embryo variability in luminal area.**

**Fig S10 Within-embryo variability in cell surface area and elongation larger than within-embryo variability in perimeter and compactness.**

**Fig S11 Morphological characteristics of non-classified endothelial cells.**

**Table S1 Frequently used mathematical symbols**.

**Table S2 Identified tuning parameter values**.

**Table S3 Robustness of chosen tuning parameter values w.r.t. cross-sectional geometry**.

**Table S4 Robustness of chosen tuning parameter values w.r.t. endothelial cell morphology**.

**Table S5 Compliance of luminal areas of estimated cross-sectional shapes with angiography**.

**Table S6 Comparison to literature-reported geometric measurements of dorsal aorta**.

**Table S7 Comparison to literature-reported morphological measurements of endothelial cells**.

## Acknowledgments

We would like to thank Anne Kühnel, Angela Hubig, Bärbel Wuntke and Julius Schwarz (University of Potsdam) for support during experiments and data acquisition. Furthermore, we would like to thank Myfanwy Evans (University of Potsdam) and Konrad Polthier and his working group (Freie Universität Berlin) for fruitful and stimulating discussions.

